# Genetic removal of Nlrp3 protects against sporadic and R345W Efemp1-induced basal laminar deposit formation

**DOI:** 10.1101/2024.10.14.618289

**Authors:** Antonio J. Ortega, Steffi Daniel, Marian Renwick, Pravallika Kambhampati, Krista N. Thompson, Gracen E. Collier, Emily L. Baker, Hasan Zaki, John D. Hulleman

**Affiliations:** Department of Ophthalmology and Visual Neurosciences, University of Minnesota, 2001 6th St. SE, Minneapolis, Minnesota, 55455, United States; Department of Ophthalmology, University of Texas Southwestern Medical Center, 5323 Harry Hines Blvd, Dallas, Texas, 75390, United States; Department of Pathology, University of Texas Southwestern Medical Center, 5323 Harry Hines Blvd, Dallas, Texas, 75390, United States

**Keywords:** chronic inflammation, inflammasome, caspase-1, AMD, EFEMP1, fibulin-3, F3, ML, DHRD

## Abstract

Chronic, unresolved inflammation has long been speculated to serve as an initiating and propagating factor in numerous neurodegenerative diseases, including a leading cause of irreversible blindness in the elderly, age-related macular degeneration (AMD). Intracellular multiprotein complexes called inflammasomes in combination with activated caspases facilitate production of pro-inflammatory cytokines such as interleukin 1 beta. Specifically, the nucleotide-binding oligomerization (NOD)-like receptor protein 3 (NLRP3) has received heightened attention due to the wide range of stimuli to which it can respond and its potential involvement in AMD. In this study, we directly tested the role of Nlrp3 and its downstream effector, caspase 1 (Casp1) in mediating early AMD-like pathology (i.e., basal laminar deposits [BLamDs]) in wild-type (WT) mice and the Malattia Leventinese/Doyne honeycomb retinal dystrophy (ML/DHRD) mouse model (p.R345W mutation in Efemp1). Compared to aged-matched controls, R345W^+/+^ knockin mice demonstrated increased Muller cell gliosis, subretinal Iba-1^+^ microglial cells, higher Nlrp3 immunoreactivity in the retina, as well as significant transcriptional upregulation of complement component 3, Nlrp3, pro-Il1b, pro-caspase-1, and tissue inhibitor of matrix metalloproteinase 3 in the retinal pigmented epithelium (RPE)/choroid. These findings were accompanied by an age-related increase in BLamD formation in the R345W^+/+^ mice. Genetic elimination of either Nlrp3 or Casp1 significantly reduced both the size and coverage of BLamDs in the R345W^+/+^ background, highlighting an important and underappreciated pathway that could affect ML/DHRD onset and progression. Moreover, Nlrp3 knockout reduced spontaneous, idiopathic BLamDs in WT mice, suggesting translatability of our findings not only to rare inherited retinal dystrophies, but also potentially to AMD itself.

## INTRODUCTION

For decades, chronic low level, unresolved inflammation has been speculated as an initiating factor in prevalent neurodegenerative diseases^1^ including Alzheimer’s disease (AD)^2^, Parkinson’s disease (PD)^3^, amyotrophic lateral sclerosis (ALS)^4^, and age-related macular degeneration (AMD)^5, 6, 7^. In many instances, this simmering level of chronic inflammation is mediated through inflammasomes, intracellular multiprotein complexes that are responsible for effecting innate immune responses through caspase-1 and the inflammatory cytokines interleukin (IL)-1β and IL-18^8^.

Depending on the stimuli, multiple distinct inflammasome complexes can form within a cell to facilitate responses to infection or stress. However, the nucleotide-binding oligomerization (NOD)-like receptor protein 3 (NLRP3) inflammasome has received heightened attention due to the wide range of stimuli to which it can respond. Canonical NLRP3 activity requires an initial transcriptional ‘priming’ step which involves either activation of toll-like receptor 4 (TLR4), the interleukin 1 receptor (IL-1R), or activation of the tumor necrosis factor receptor (TNFR)^9^. These signals, which are also known as pathogen associated molecular patterns (PAMPs), in turn promote the nuclear factor kappa B (NF-κB)-dependent transcription of critical components of the NLRP3 inflammasome, including NLRP3 itself, pro-caspase-1, and pro-IL-1β^9, 10^. A second ‘activating’ signal, or damage/danger-associated molecular pattern (DAMP), is typically required for full NLRP3 inflammasome activity, which results in the processing of pro-caspase-1 (CASP1) followed by cleavage of pro-IL-1β and pro-IL-18 and their secretion^11^. Such DAMPs or secondary signals can originate from a wide variety of sources including K^+^ efflux, mitochondrial dysfunction/oxidative stress, glucose, lysosomal rupture, extracellular ATP, complement activation, aggregated protein such as amyloid beta, or even cholesterol crystals^1^. While the two-step NLRP3 activation mechanism has received the most focus and attention, groups have also demonstrated one-step NLRP3 activation through TLR4 stimulation in human and porcine monocytes^12, 13^, and simultaneous activation of TLRs along with the presence of ATP can trigger rapid NLRP3 activation^14^. Interestingly, whereas it is widely accepted that NLRP3 is primarily expressed in immune cells (e.g., monocytes and macrophages), multiple studies suggest that very low levels of NLRP3 may be present in a variety of tissues^15^ including neurons^16^, islets^17^, and retinal cells^18, 19^, providing a localized response to infection and/or stress.

The ultimate result of full activation of the NLRP3 inflammasome (and subsequent CASP1 activation) is to produce and secrete pro-inflammatory cytokines including IL-1β and IL-18, which can serve as recruitment signals for immune cells and initiate clearance of viruses, pathogens, or microbes^20^. Additionally, canonical or non-canonical inflammasome activation can potentially lead to CASP1-triggered gasdermin D-dependent cell lysis or ‘pyropoptosis’^9^, which, in turn, can release additional pro-inflammatory intracellular antigens at the site of stress or infection^21^. Accordingly, activation of the NLRP3 inflammasome is critical for remediation of acute viral or microbial infections^22^. Yet, if persistently activated, sustained NLRP3 activity can cause chronic and unresolved inflammation, leading to tissue damage and degeneration of sensitive cell lineages, including neurons^23^. Thus, reducing chronic NLRP3 inflammasome activation has been a prime target for minimizing age-related neurodegenerative pathology in both the brain and eye^19, 24, 25, 26, 27^.

AMD is a progressive, age-related degenerative ocular disease that affects the macula, the central portion of the retina containing cone photoreceptors responsible for high-acuity vision, such as when an individual reads or recognizes faces^28^. AMD is the leading cause of irreversible blindness in the elderly in industrialized nations and is predicted to affect almost 300 million people over the age of 55 worldwide by 2040^29^. Patients with AMD are characterized by the development of extracellular lipid and protein-rich deposits called ‘drusen’ underneath a critical photoreceptor support cell, the retinal pigmented epithelium (RPE)^30^. As drusen coalesce and become ‘softer’ in appearance and composition, the risk for compromising the RPE and developing intermediate or advanced (blinding) forms of AMD increases. Unfortunately, the molecular causes of AMD are still not completely understood, but likely involve chronic inflammation, altered lipid metabolism, increased oxidative stress, and impaired extracellular matrix (ECM) maintenance^31, 32^. Interestingly, the NLRP3 inflammasome links all four of these potential AMD contributors^33, 34, 35, 36^. Indeed, a series of studies have implicated NLPR3 or its downstream products in the progression of different stages of AMD. Whereas the Ambati group found a detrimental role for the NLRP3 inflammasome in the onset of geographic atrophy (GA, i.e., loss of RPE cells)^19, 37, 38, 39^, the Cambell group found that downstream cytokines such as IL-18 appear to protect against neovascularization of the retina (i.e., wet AMD)^40, 41, 42, 43^. More recently, a series of studies have demonstrated a detrimental role of NLRP3 in additional, non-AMD degenerative retinal diseases^44, 45^. Further supporting the notion of role of NLRP3 in pathology resembling AMD, recently a 38 yo Chinese female with an autosomal dominant, autoimmune mutation in NLRP3 (c.1043C>T, p.T348M) presented with pseudodrusen (amorphous material between the RPE and photoreceptors) and hard drusen near the optic nerve^46^. These additional pieces of evidence strongly suggest a pathogenic role of chronic NLRP3 activation in the eye.

Yet, as an etiologically complex and primate-specific age-related disease, it is difficult to study and confirm the molecular underpinnings of AMD within a laboratory and on a reasonable timeframe. To overcome this challenge, we^47, 48, 49, 50, 51, 52, 53, 54, 55, 56, 57^ and other groups^58, 59, 60, 61, 62, 63, 64, 65, 66, 67, 68, 69, 70, 71, 72^ have focused our attention on a monogenic, autosomal dominant, early-onset form of AMD called Malattia Leventinese/Doyne honeycomb retinal dystrophy (ML/DHRD). ML/DHRD is a rare macular dystrophy that is caused by a p.R345W mutation in the extracellular matrix protein, epidermal growth factor-containing fibulin-like extracellular matrix protein 1 (EFEMP1)^73^, also known as fibulin-3 (FBLN3). Although it is more common for individuals with this mutation to develop disrupted vision in their third or fourth decade^74^, in the most severe documented case, a male child who was homozygous for the R345W mutation, developed extensive drusen deposition and RPE atrophy by age 12^75^. While the NLRP3 inflammasome has not been directly implicated in the pathology of ML/DHRD to date, extensive findings in mice and cultured cells implicate a key role for immune-related complement activation in R345W BLamD pathology^69, 70, 71, 72, 76, 77, 78, 79^, a clinically-observed pathology that likely contributes toward the formation/progression of high-risk drusen in AMD patients^80^. Moreover, primary mouse RPE cell cultures expressing R345W produced elevated levels of Il1β and Il6^70^. Neutralizing these pro-inflammatory cytokines prevented the formation of F3-dependent amorphous deposits found in culture^70^. The culmination of these observations suggest the involvement of the NLRP3/Nlrp3 inflammasome in the progression of ML/DHRD. Herein, we directly tested this possibility through the generation and ocular characterization of R345W^+/+^ knockin mice lacking either Nlrp3 or Casp1. Whole body genetic removal of either of these inflammasome components significantly reduced the size and coverage of the primary phenotype in ML/DHRD mice, BLamDs. Moreover, Nlrp3 knockout reduced the size and coverage of spontaneous BLamDs in a WT C57BL/6 background, suggesting translatability of our findings not only to rare inherited retinal dystrophies (ML/DHRD), but also potentially to sporadic AMD.

## MATERIALS AND METHODS

### Animal housing, handling and approval

All mice used in these studies were either obtained directly from Jackson Laboratories (JAX, Bar Harbor, ME) or produced in house. Mice were housed under 12 h on/12 h off light cycles and fed standard rodent chow/water ad libitum at either UT Southwestern (Teklad Global 2016 Rodent Diet) or the University of Minnesota (UMN, Teklad Global 2918 Rodent Diet). All procedures were conducted under an Institutional Animal Care and Use Committee (IACUC)-approved animal protocol (#2016-101740 at UT Southwestern, #2308-41350A at UMN) and followed the ARVO guidelines for the "Use of Animals in Ophthalmic and Vision Research”. All mice analyzed were on the C57BL/6 background (or C57BL/6J if directly from Jackson Laboratories). Malattia Leventinese/Doyne honeycomb retinal dystrophy (ML/DHRD) mice harboring two copies of the p.R345W Efemp1 mutation (R345W^+/+^) were obtained from Harvard Medical School (Boston, MA, courtesy of Dr. Eric Pierce) and were characterized previously^75^. Nlrp3 knockout (Nlrp3^-/-^) mice (JAX stock #021302^13^) and caspase 1/11 knockout (Casp1^-/-^) mice (JAX stock # 016621^81^) were provided by Hasan Zaki (UT Southwestern, Dallas, TX) but originated from JAX. All breeder mice were confirmed to not harbor common retinal degeneration mutations (e.g., *rd8* and *GNAT*). Sex as a biological variable was taken into account, but we did not detect any sex-based differences in our studies.

### Spectral domain optical coherence tomography (SD-OCT)

Twenty-month-old WT and R345W^+/+^ mice were anesthetized by an i.p. ketamine/xylazine injection followed by pupil dilation (1% tropicamide, Sandoz, Switzerland) prior to obtaining B-scans using an Envisu R2210 SD-OCT (Bioptigen, Morrisville, NC). Parameters were as follows: depth - 1.65 mm, width – 1.8 mm, depth samples – 1024, width samples – 1000, length samples – 1000, depth pitch – 1.61 um/pix, width pitch – 1.80 um/pix, length pitch – 1.80 um/pix, refractive index – 1.38, rectangular scan type. B-scans were then exported as high-resolution TIFF images. Automated retinal layer segmentation and individual layer thickness measurements were performed using the InVivoVue Diver software (Bioptigen). Results were then exported and analyzed using a two-way ANOVA (Prism, San Diego, CA).

### Neural retina/RPE flat-mounts

Enucleated eyes were post-fixed in 4% paraformaldehyde (PFA, Electron Microscopy Sciences, Hatfield, PA) for 2 h at RT, followed by washes (3x 10 min) in phosphate buffered saline (PBS). Posterior eye cups were dissected and the neural retina was separated from the RPE/choroid. Once separated, they were pre-treated with 0.3% triton X-100 in PBS (4x 30 min), and subsequently blocked for 2 h in 0.1% triton X-100 (Fisher Scientific, Pittsburgh, PA) in PBS containing 10% normal goat serum (New Zealand origin, Gibco, Waltham, MA) with 1% bovine serum albumin (Fisher Bioreagents, Waltham, MA). Retinal ganglion cells (RGCs) in the neural retina were labeled using an anti-RNA-Binding Protein with Multiple Splicing (RBPMS, rabbit IgG, 1:500, GeneTex, #GTX118619, Irvine, CA) antibody, microglia in the neural retina and RPE were labeled using anti-ionized calcium-binding adaptor molecule 1 (Iba1, rabbit IgG, 1:200, FUJIFILM Wako Chemicals, Richmond, VA) antibody, and tight junctions in the RPE were labeled using anti-zonula occludens-1 (ZO-1, rabbit IgG, 1:250, Invitrogen, #40-2200, Carlsbad, CA) antibody overnight at 4°C. After PBS washes, the neural retina/RPE were incubated overnight at 4°C with goat anti-rabbit AlexaFluor488 secondary antibody (1:1000, Life Technologies, #A27034, Carlsbad, CA) or goat anti-rabbit AlexaFluor 594 secondary antibody (1:1000, Invitrogen #A-11012, Carlsbad, CA)) diluted in 0.1% Triton X-100 in PBS and then mounted with Vectashield Mounting Medium containing 4′,6-diamidino-2-phenylindole (DAPI, Vector Laboratories, Burlingame, CA). Confocal microscopy (performed on a Leica TCS SP8 or Leica Stellaris 8) was used to capture three images each from the peripheral, mid-peripheral, and central regions of the four quadrants of each neural retina and RPE for quantification studies (12 images per retina) at 20/25x magnification. RGC counts were performed using the ImageJ (FIJI) Cell Counter Plugin as described previously^55^. For Iba1 quantification, image stacks were processed in FIJI and analyzed using the MorphoLibJ plugin. ZO-1 stained RPE flat-mounts were imaged in stacks at 20x magnification (on a Nikon AX R with NSPARC) then maximum projected and tiled to generate one single image per eye. These were further processed for morphometric analyses using the Retinal Epithelium Shape and Pigment Evaluator (REShAPE) platform (see below)^82, 83^.

### Retinal pigment epithelium (RPE) morphometry analysis

Retinal Epithelium Shape and Pigment Evaluator (REShAPE)^83^, an advanced artificial intelligence-based software, was used to analyze the morphometric properties of RPE cells in whole-mounted eye specimens from 20-month-old mice. A total of 13 RPE flatmounts were examined, comprising six WT mice and seven R345W^+/+^ mice. REShAPE facilitated comprehensive analysis by providing segmented images and detailed morphometric data files for individual cells. The software also generated color-coded heatmaps reflecting various morphological features, including cell area, aspect ratio, hexagonality, and the number of neighboring cells. The automation of RPE subpopulation analysis was achieved through Python and batch processing with macros in Fiji/ImageJ. The cell area heatmaps were segregated into three concentric subpopulations within each RPE flatmount, designated as P1, P2, and P3, progressing from the innermost to the outermost regions. These subpopulations were subsequently isolated and REShAPE was used to conduct a detailed morphometric analysis of each subpopulation individually.

### Eye orientation for histology and transmission electron microscopy (TEM)

Previous studies have demonstrated an enrichment of BLamDs in the ventral-temporal quadrant of the mouse retina^76^. To increase consistency and comparability across samples, we analyzed BLamDs only within this quadrant. Prior to enucleation, eyes were marked temporally with a Bovie cautery fine tip. Post fixation, an incision was made at this same mark and the eye cut along the ora serata leaving behind a corneal flap overlaying the quadrant of interest.

### Sample processing for histology and TEM

Four to twenty-month-old WT or R345W^+/+^ mice were deeply anesthetized using an i.p. ketamine/xylazine injection. Additionally, Casp1^-/-^ R345W^+/+^ and Nlrp3^-/-^ R345W^+/+^ mice were also aged to sixteen-months and their eyes harvested. Briefly, once unresponsive to foot pinch, mice were immediately transcardially perfused with saline followed by 10 mL of 4% v/v paraformaldehyde (PFA, Electron Microscopy Sciences, Hatfield, PA). Prior to enucleation, an incineration mark was made (described above) to orient the eye. Eyes for histology were enucleated and processed for freeze substitution as described previously^84, 85, 86^ and submitted to the UT Southwestern Histo Pathology Core. Sections were taken through the optic nerve and images were recorded on a NanoZoomer S60 microscope (Hamamatsu, Bridgewater, NJ).

The contralateral eye from each mouse was processed for TEM analysis by placing it in half strength Karnovsky’s fixative (2% v/v formaldehyde + 2.5% v/v glutaraldehyde, in 0.1M sodium cacodylate buffer, pH 7.2) on a rocker at RT for 4 h. An incision was made along the ora serrata removing the lens and majority of the cornea, while using the temporal incineration mark to leave a corneal flap over the ventral temporal quadrant of interest. The eye cups were placed in fresh half strength Karnovsky’s fixative overnight at 4°C. Eye cups were prepared for TEM by rinsing three times in 0.1M sodium cacodylate buffer followed by 1% v/v OsO_4_ in buffer (2 parts 0.1M sodium cacodylate buffer pH 7.2, 1 part 4% w/v osmium, 1 part 6% v/v potassium ferrocyanide) for 2 h at RT and washed three times in 0.1M sodium cacodylate buffer (pH 7.2) for 30 min at RT. The sample was then rinsed twice with Nanopure water, placed in 2.5% uranyl acetate dissolved in Nanopure water for 20 min followed by 25% ethanol for 30 min, 50% ethanol for 30 min, 75% ethanol for 30 min, 95% ethanol for 45 min, then two serial 100% ethanol incubations for 45 min each (all steps performed at RT) and kept overnight at 4°C.

Samples were then resin infiltrated by first replacing the ethanol by performing two 100% propylene oxide washes for 20 min, then 2 parts 100% propylene oxide per 1 part Eponate 12 resin (Ted Pella, Redding, CA) for 1 h, 1 part 100% propylene oxide per 2 parts Eponate 12 resin for 1 h, then left overnight in 100% Eponate 12 (all steps performed at RT). The next day samples were placed in fresh Eponate 12 for 2 h then embedded with in Eponate 12 resin in a mold and placed in a 60°C oven overnight. Samples were cut using the corneal flap as an orientation marker such that sections were obtained from the ventral temporal region. Sections mounted grids were stained with uranyl acetate and lead citrate contrast. The grids were imaged at 1,200x magnification using a 1400+ TEM (JEOL, Akishima, Japan). TIFF file images were acquired using the interfaced BioSprint ActiveVu mid-mount CCD camera (Advanced Microscopy Technologies, Woburn, MA).

TEM images were analyzed with Fiji software. Brightness and contrast were adjusted for best visualization of Bruch’s membrane (BrM). Images were excluded from analysis if the membrane and/or deposits could not be reasonably delineated. Prior to tracing, pixels were scaled to µm. For BLamD size analysis, BLamDs and the RPE basement membrane were traced in deposit containing fields of view (FOVs), using the Fiji polygon selection tool. The areas of the traced deposits and membranes were reported in µm^2^. For BLamD coverage analysis, FOVs were divided into equal sub-FOVs, along the BrM, measuring 2.23 µm in length using the grid function. Masked observers determined whether deposits were present in each sub-FOV with a binary classification (100 = deposit, 0 = no deposit). Data are reported as percent BLAMD coverage. The presence of deposits was additionally verified by a secondary observer. Once the images were analyzed, either Kruskal-Wallis or Mann-Whitney tests were performed comparing genotypes across ages, with p < 0.05 as the cutoff for significance.

### Quantitative PCR (qPCR) analysis

Mice were anesthetized using a ketamine/xylazine cocktail via i.p. injection. Eyes were enucleated and dissected to remove the anterior segment and the posterior eye cup was further dissected to obtain neural retina and RPE/choroid. RNA isolation was carried out using the Aurum Total RNA isolation kit (BioRad, Hercules, CA). Samples were reverse-transcribed to cDNA using qScript cDNA Supermix (Quantabio, Beverly, MA). Available TaqMan probes (Applied Biosystems, Waltham, MA) were used for qPCR reactions as follows: β-actin (Mm02619580_g1), C3 (Mm01232779_m1), Casp1 (mm00438023_m1), Cfh (Mm01299248_m1), Efemp-1 (Mm00524588_m1), Gfap (Mm01253033_m1), Il18 (mm00434226_m1), Il1β (Mm00434228_m1), Il6 (Mm00446190_m1), Nlrp3 (mm00840904_m1), and Timp3 (Mm00441826_m1). Samples were run on a QuantStudio 6 Real-Time PCR system (Applied Biosystems) in technical and biological replicates. Relative abundance was calculated by comparing expression to β-actin using the QuantStudio software.

### Immunohistochemistry

Unstained paraffin embedded sections (Histo Pathology Core, UT Southwestern), were de-paraffinized, and subjected to antigen retrieval using the EZ-Retriever System (BioGenex, Fremont, CA). Slides were washed (2x 5 min) in 1x Tris-buffered saline (TBS) followed by pretreatment in 0.025% Triton X-100/1xTBS with gentle agitation (2x 5 min). Sections were blocked in blocking solution consisting of 10% normal goat serum (New Zealand origin, Gibco, Waltham, MA) with 1% bovine serum albumin (Fisher Bioreagents, Waltham, MA) in TBS for 2 h at RT in a humidified chamber. Slides were drained and incubated in primary antibodies diluted in blocking solution overnight at 4 °C. The antibodies used were as follows: anti-NLRP3 antibody (rabbit IgG, 1:200, Cell Signaling, #15101S, Danvers, MA), and anti-glial fibrillary acidic protein (GFAP) antibody (rat IgG2a, 1:250, Invitrogen, #13-0300, Carlsbad, CA). The next day, slides were rinsed with 0.025% triton X-100 in TBS (2 × 5 min) and incubated in goat anti-rabbit AlexaFluor488 secondary antibody (1:1000, Life Technologies, #A27034, Carlsbad, CA) or goat anti-rat AlexaFluor594 secondary antibody (1:1000, Life Technologies, #A48264) in 0.025% triton X-100 in TBS overnight at 4 °C. Vimentin was labeled using Vimentin-Label Atto488 (1:200, ChromoTek, #vba448, Rosemont, IL) overnight at 4 °C. Sections were then stained with 4′,6-diamidino-2-phenylindol (DAPI) at RT for 20 min, rinsed with TBS (3 × 5 min) and mounted with Vectashield (Vector Laboratories, Newark, CA). Images were taken using a 25x or 63x objective on a TCS SP8 or Stellaris 8 microscope (Leica Microsystems, Buffalo Grove, IL).

## RESULTS

### R345W^+/+^ mice retain gross retinal and RPE structure even during advanced age

R345W^+/+^ mice serve as an essential in vivo model system for studying the molecular mechanisms underlying ML/DHRD and potential treatments for this rare disease. Previous studies have indicated that R345W^+/+^ mice develop age- and genotype-dependent accumulation of BLamDs^67, 75^, accumulations of lipids, apolipoproteins, and extracellular matrix proteins including fibronectin^87^, laminin, collagens^88^, and EFEMP1^67, 75^ within the basal lamina of RPE. Yet, whether these deposits lead to retinal thinning and degeneration in mice is less clear. We therefore began our studies by performing spectral domain optical coherence tomography (SD-OCT) on aged WT and R345W^+/+^ mice at advanced age (20 mo) followed by automated quantification of retinal layer thickness (Fig. 1A-C). Interestingly, WT and R345W^+/+^ mice showed nearly identical total retinal thickness (225.85 ± 5.42 μm and 227.68 ± 8.18 μm, respectively) as well as similar thickness of individual retinal cell layers (Fig. 1A-C), findings that were also validated using standard hematoxylin and eosin (H&E) histology of the retina (Fig. 1D, E).

**Fig. 1.**
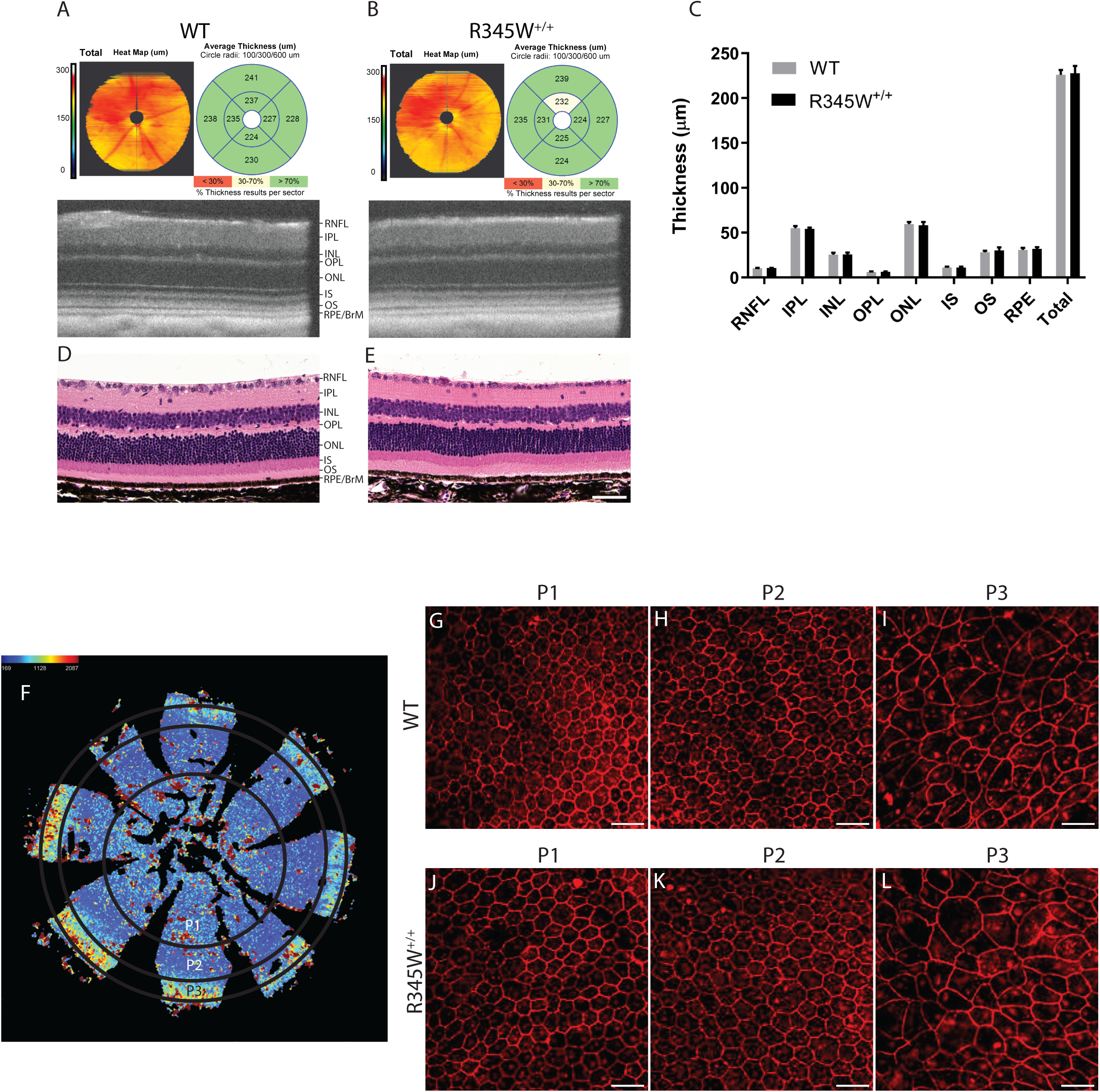
No gross structural changes in the retina or RPE are observed in WT vs. R345W^+/+^ mice even at ≥20 months of age. (A, B) Spectral domain optical coherence tomography (SD-OCT) en face and B-scans of 20 mo WT and R345W^+/+^ mice. Representative images of n > 20 mice. (C) Quantification of retinal layers using Diver software. n for WT = 28 eyes (20 male, 8 female), n for R345W^+/+^ = 21 (12 males, 9 females). Retinal nerve fiber layer (RNF), inner plexiform layer (IPL), inner nuclear layer (INL), outer plexiform later (OPL), outer nuclear layer (ONL), inner segment (IS), outer segments (OS), and retinal pigmented epithelial layer/Bruch’s membrane (RPE/BrM). Mean ± S.D. presented. No values were significant as determined by a two-way ANOVA multiple comparisons test. (D, E) H&E histology of WT and R345W^+/+^ retinas at 20 mo of age. Representative images of n=>3 mice. Scale bar = 50 μm. (F) Illustration of an RPE flatmount segmented into different RPE subpopulations based on morphometric parameters. (G-I) Representative images of P1-P3 subpopulations in 22 mo WT mice (n = 6). (J-L) Representative images of P1-P3 subpopulations in 22 mo R345W^+/+^ mice (n = 7).

Next, we examined more closely RPE size, hexagonality, aspect ratio, and neighbor number using flatmounts (Fig. 1F-L) followed by systematic quantification of these parameters using REShAPE in a similar manner as performed previously using human RPE flatmounts^82, 83^. Our findings indicated that in aged mice (22 mo), the P3 subpopulation, located in the far peripheral region (Fig. 1F), exhibited the largest cell area, followed by the innermost P1 and then P2 subpopulations (Fig. 1I, L, Sup. Fig. 1A). Furthermore, there was a notable increase in cell elongation from the innermost to the outermost regions, accompanied by a corresponding decrease in hexagonality (Fig. 1G-L, Sup. Fig. 1B-D). However, when these parameters were compared between the two genotypes, no statistically significant differences were observed among any of the subpopulations (Sup. Fig. 1A-D), indicating similar RPE morphology (and likely functionality) between WT and R345W^+/+^ mice.

Given the recent findings that mutations in EFEMP1 can cause juvenile onset glaucoma^89, 90, 91^, and that the R345W mutation may predispose some human patients to corneal opacity^92^ or even glaucoma itself^93^, we tested whether the R345W mutation predisposed mice to corneal changes, elevated intraocular pressure (IOP) or retinal ganglion cell (RGC) death. Corneas from 14 mo WT and R345W^+/+^ mice were unremarkable by slit lamp examination (Sup. Fig. 2A), in contrast to our previous observations in aged Efemp1 KO mice that demonstrated age-dependent corneal opacity and vascularization by this age^56^. Conscious, daytime IOPs evaluated using a rebound tonometer were not significantly different between WT and R345W^+/+^ mice (14 mo, Sup. Fig. 2B). Moreover, RGC number as determined by RNA-binding protein with multiple splicing (RBPMS) staining and quantification was also not significantly different between the two genotypes, even at 17 mo (Sup. Fig. 2C, D). These results suggest that the R345W mutation has no detectable effects on anterior chamber biology in mice.

Finally, we also assessed whether the R345W mutation predisposed mice to systemic body-wide metabolic changes including elevated cholesterol, triglycerides, or even body composition that might explain their apparent predisposition to lipid-rich BLamD formation^67, 75^. However, we did not detect any difference in circulating cholesterol (Sup. Fig. 3A), triglycerides (Sup. Fig. 3B), body weight, or composition (Sup. Fig. 3C, aside from gender differences in total weight, favoring heavier males) between WT and R345W^+/+^ mice.

### Quantitative longitudinal transmission electron microscopy (TEM) highlights natural history disease progression of ML/DHRD in mice and reveals critical windows of disease advancement

While macroscopically the WT and R345W^+/+^ mice appeared similar, we next focused on potential ultrastructural differences at the RPE/BrM interface in the form of BLamDs. Due to their relatively small size, BLamD quantification in mice usually requires TEM for detection and quantification. Previously, research groups have typically provided representative TEM FOVs for illustrative purposes. However, when we analyzed WT and R345W^+/+^ TEM images, we noticed that there was substantial variability across FOVs with respect to BLamD prevalence and size (typically observed in WT mice < 16 mo and R345W^+/+^ < 8 mo). Therefore, we devised a quantitation strategy for BLamD measurement and undertook a thorough study of nearly 1000 FOVs in WT and R345W^+/+^ mice (Fig. 2A-D) spanning 4 - 20 mo, quantifying BLamD size and coverage in a masked approach. Wild-type mice formed few periodic small BLamD deposits between 4 mo and 16 mo (Fig. 2A, C, Sup. Fig. 4A). However, during this time, the number of FOVs that contained deposits rather consistently increased at each timepoint (Fig. 2A, D, Sup. Fig. 4B). Interestingly, with advanced age (i.e., 20 mo), spontaneous BLamDs in WT mice continued to form and we observed a large and significant jump in BLamD coverage in mice between 16 and 20 mo (Fig. 2A, C, D, Sup. Fig. 4A, B, **** p < 0.0001). Like WT mice, R345W^+/+^ mice also did not form large BLamD early in their lifespan between 4 and 8 mo (Fig. 2B, C, Sup. Fig. 4C). Surprisingly, R345W^+/+^ mice had slightly (but significantly) lower extents of BLamD coverage when compared to WT mice at 8 mo (Fig. 2B, D, Sup. Fig. 4D, * p < 0.05). However, R345W^+/+^ mice demonstrated an explosive and significant jump in BLamD size at a relatively young age (8 to 12 mo, Fig. 2B, C, Sup. Fig. 4C, **** p < 0.0001) which subsequently plateaued for the remaining timepoints (Fig. 2B, C, Sup. Fig. 4C). During each of the timepoints taken, R345W^+/+^-induced BLamD coverage steadily increased, reaching almost 60% of the sub-FOVs analyzed (Fig. 2B, D, Sup. Fig. 4D). Quite surprisingly however, by 20 mo of age, WT and R345W^+/+^ mice were indistinguishable with respect to BLamD coverage (Fig. 2A, B, D) although R345W^+/+^ BLamDs were significantly larger (on average ∼2.9 times) than WT BLamDs (Fig. 2A-C, **** p < 0.0001).

**Fig. 2.**
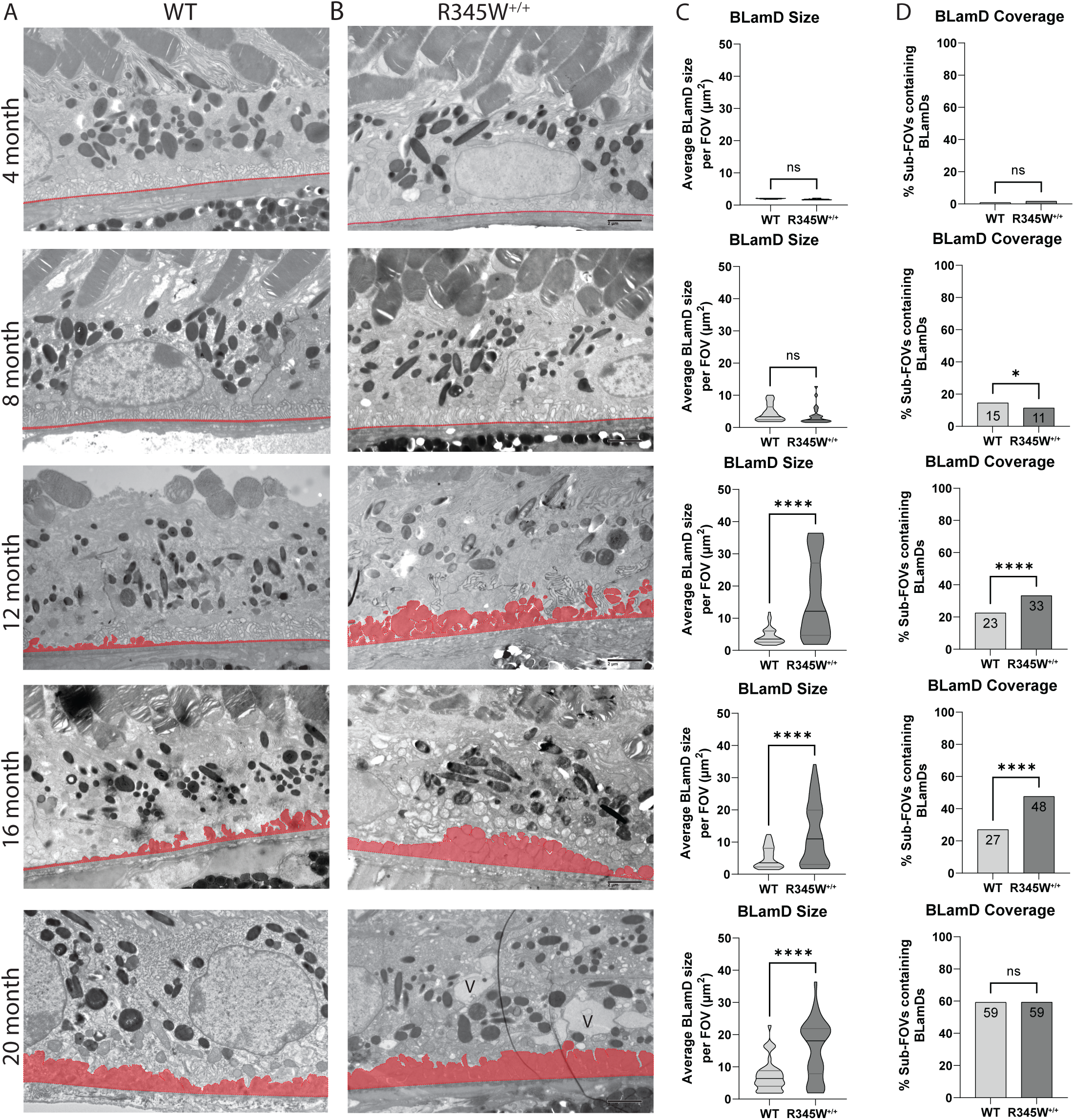
Quantitative, longitudinal transmission electron microscopy (TEM) delineates basal laminar deposit (BLamD) progression in WT and R345W^+/+^ mice. (A, B) TEM analysis of WT (A) or R345W^+/+^ (B) mice at 4 mo intervals ranging from 4-20 mo. Representative images of ∼16 FOVs per mouse, n = 6 mice/genotype and age group (3 male, 3 female) except for 20 mo mice. n = 3 females/genotype at 20 mo. Scale bar = 2 μm. Area highlighted in red designates the tracing used to quantify BrM and BLamDs (if present). Images taken at 2,500x magnification. Scale bar = 2 μm. V = vacuole. (C, D) Quantitation of BLamD size (µm^2^/FOV) (C) and percent sub-FOVs (D) that contain at least one BLamD (see Materials and Methods for in-depth information on definition and quantitation of BLamDs). Data in (C) are presented as violin plots showing the median and quartiles. Data in (D) are presented as percent of total FOVs counted for each age/genotype. Comparative analysis performed using Mann-Whitney tests. * p < 0.05, **** p < 0.0001. Note: The WT TEM image and WT BLamD quantification are identical to that shown later in Fig. 10B-D.

### Comparative molecular analysis points to the involvement of chronic inflammation and activation of the Nlrp3 inflammasome in aged R345W^+/+^ mice

While we did not note any macroscopic changes to the gross structure of the retina or RPE in aged R345W^+/+^ mice, given such notable ultrastructural BLamD pathology, we began to ask whether the R345W^+/+^ RPE was reacting to, or being modified molecularly as a consequence of the Efemp1 protein mutation or BLamD formation. Previous groups have implicated the R345W mutation as a trigger of inflammation, complement activation, and membrane attack complex deposition in vivo and in cells^69, 70, 71, 72^. Thus, we decided to assess whether we could transcriptionally detect elevated complement and inflammatory markers in the RPE of aged (18 mo) R345W^+/+^ mice. Consistent with previous findings, we detected elevated transcripts of complement component 3 (C3), and interleukin 1 beta (Il1β) in the RPE/choroid of R345W^+/+^ mice (Fig. 3A, * p < 0.05). Additionally, we noted significant increases in inflammasome components, caspase-1 (Casp1) and NOD-like receptor protein 3 (Nlrp3) in RPE/choroid samples (Fig. 3A, * p < 0.05, *** p < 0.001), corroborating a general state of inflammation in R345W^+/+^ RPE/choroid at advanced age.

**Fig 3.**
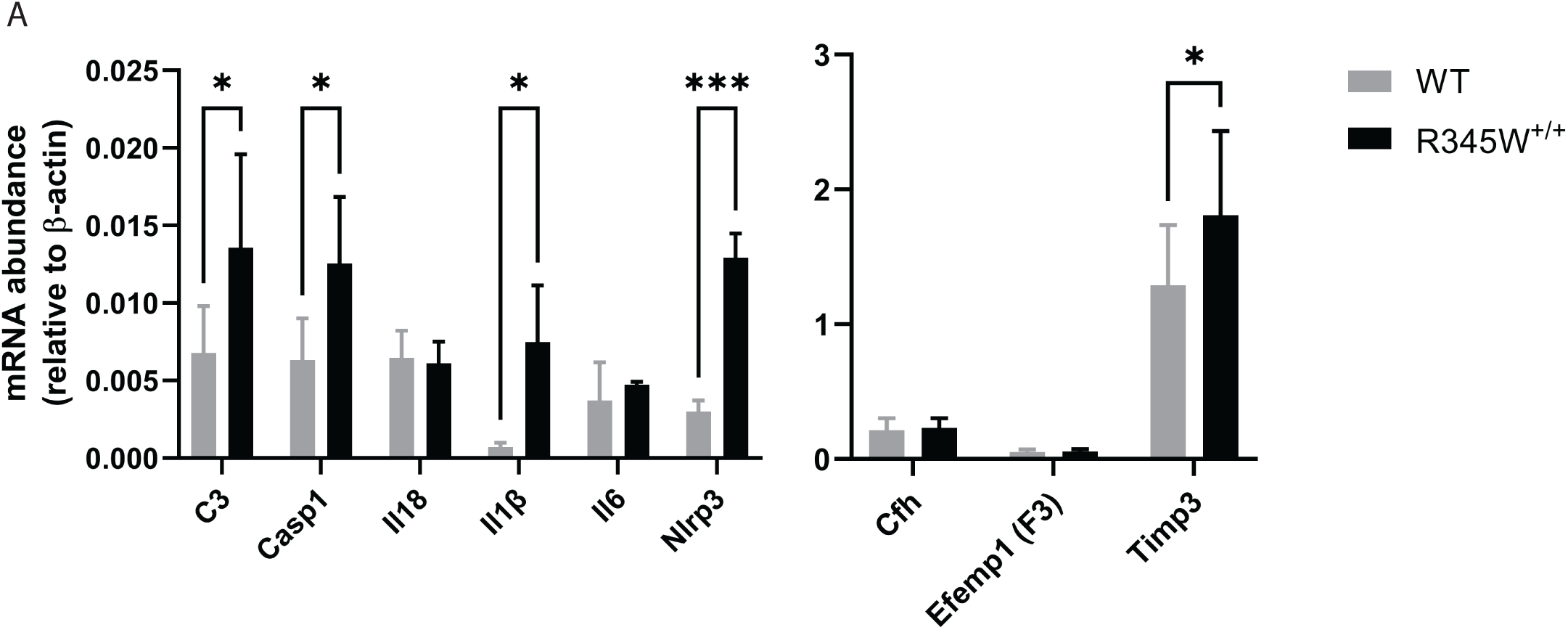
Transcriptional analysis suggests a state of chronic inflammation in R345W^+/+^ RPE/choroid at 18 mo. (A) mRNA was extracted from WT or R345W^+/+^ RPE/choroid samples, reverse transcribed and probed for transcript levels of complement component 3 (C3, p ≤ 0.05), caspase 1 (Casp1), interleukin 18 (Il18), interleukin 1 beta (Il1β), interleukin 6 (Il6), nucleotide-binding oligomerization (NOD)-like receptor protein 3 (Nlrp3), complement factor H (Cfh), Efemp1 (F3), tissue inhibitor of matrix metalloproteinase 3 (Timp3) and plotted relative to β-actin (n = 5-6 mice). 2-way ANOVA. Mean ± S.D. * p < 0.05.

Next, we confirmed upregulation of Nlrp3 in the retina of R345W^+/+^ mice using retinal immunohistochemistry across a series of ages ranging from 4-20 mo (Fig. 4A-H). At young ages (4-8 mo), we observed virtually no Nlrp3 staining in the retina in either genotype (Fig. 4A, B, E, F). However, beginning at 12 mo (when BLamD size peaked in R345W^+/+^ mice, Fig. 2C), we detected a noticeable increase in immunoreactivity for Nlrp3 that was greater than in WT mice (Fig. 4C, G). This increase in Nlrp3 staining continued to grow at 20 mo, creating an even more striking difference in Nlrp3 signal between WT and R345W^+/+^ mice (Fig. 4D, H). Nlrp3^-/-^ mice were used to demonstrate antibody specificity (Fig. 4I).

**Fig 4.**
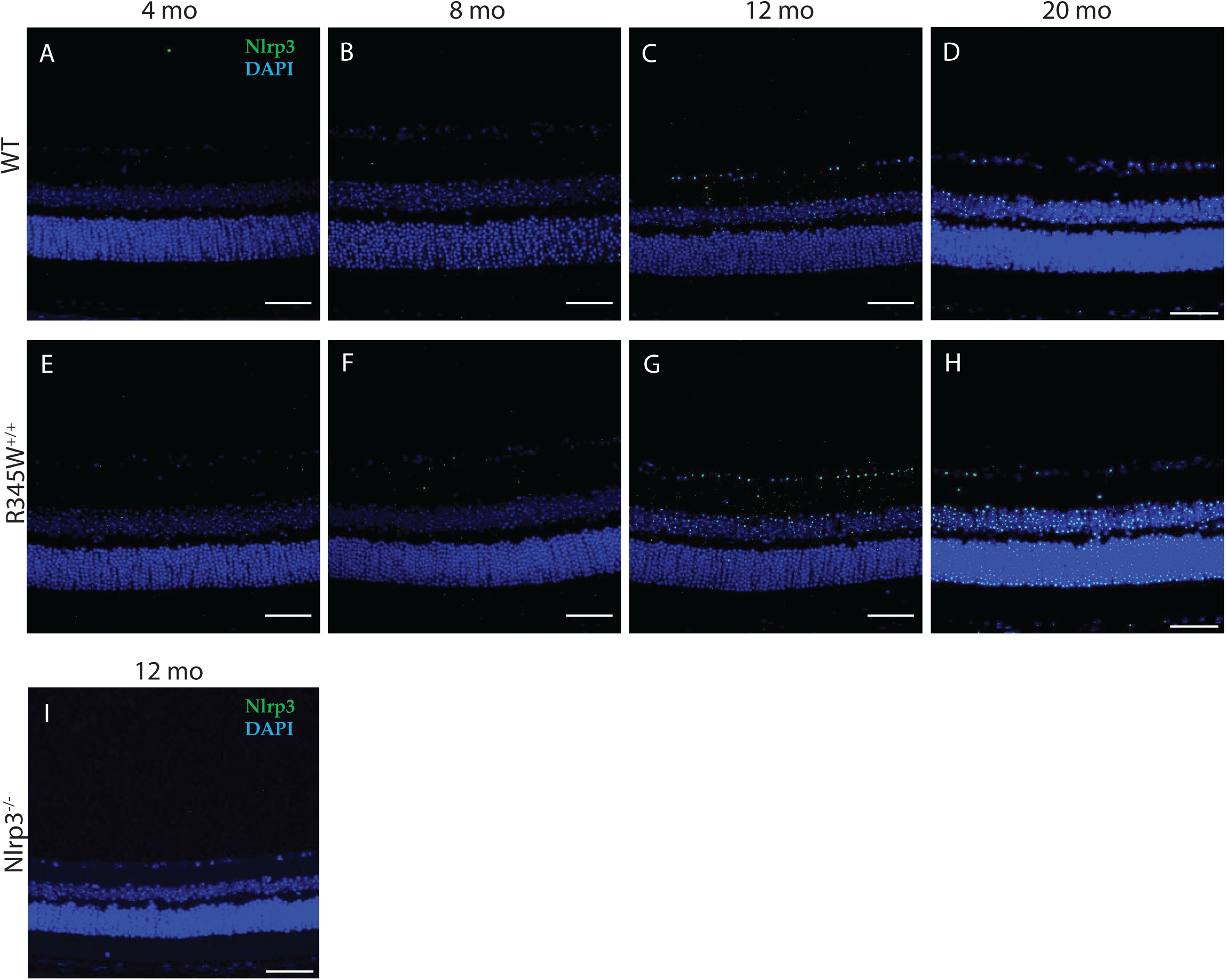
Increased Nlrp3 expression observed in aging R345W^+/+^ mouse retina compared to WT mice. (A-H) Representative images of retinal cross sections showing expression of NLRP3 (green) along with nuclear stain DAPI (blue) (scale bar = 50 µm) across ages (A, E) 4 mo, (B, F) 8 mo, (C, G) 12 mo, (D, H) 20 mo. Elevated Nlrp3 inflammasome immunoreactivity is observed in aging R345W^+/+^ mice compared to their WT littermates beginning at 12 mo of age. (I) No detectible Nlrp3 staining in Nlrp3 deficient mice.

### Increased Muller cell gliosis and activated Iba1^+^ microglia are observed in aged R345W^+/+^ <u>mouse retina</u>

Muller cells are the most common glial cell found in the retina and support a wide variety of structural and functional roles ranging from metabolism to neurotransmitter regulation to retinal cell regeneration^94^. Given the Muller cell’s exquisite ability to sense retinal stretch^95^, we asked whether the BLamDs generated by the R345W mutation may result in Muller cell gliosis. To test this hypothesis, we probed for Vim and Gfap retinal staining across mouse ages of 4-19 mo in both the WT and R345W^+/+^ genotypes (Fig. 5A-P). Beginning at 8 mo, we observed prominent Vim and Gfap staining in R345W^+/+^ mice (Fig. 5F, N) near the RGC layer and IPL that was not present in WT counterparts (Fig. 5B, J). This increased Vim and Gfap staining persisted, unresolved for the remainder of the life of the R345W^+/+^ mouse (Fig. 5G, H, O, P).

**Fig. 5.**
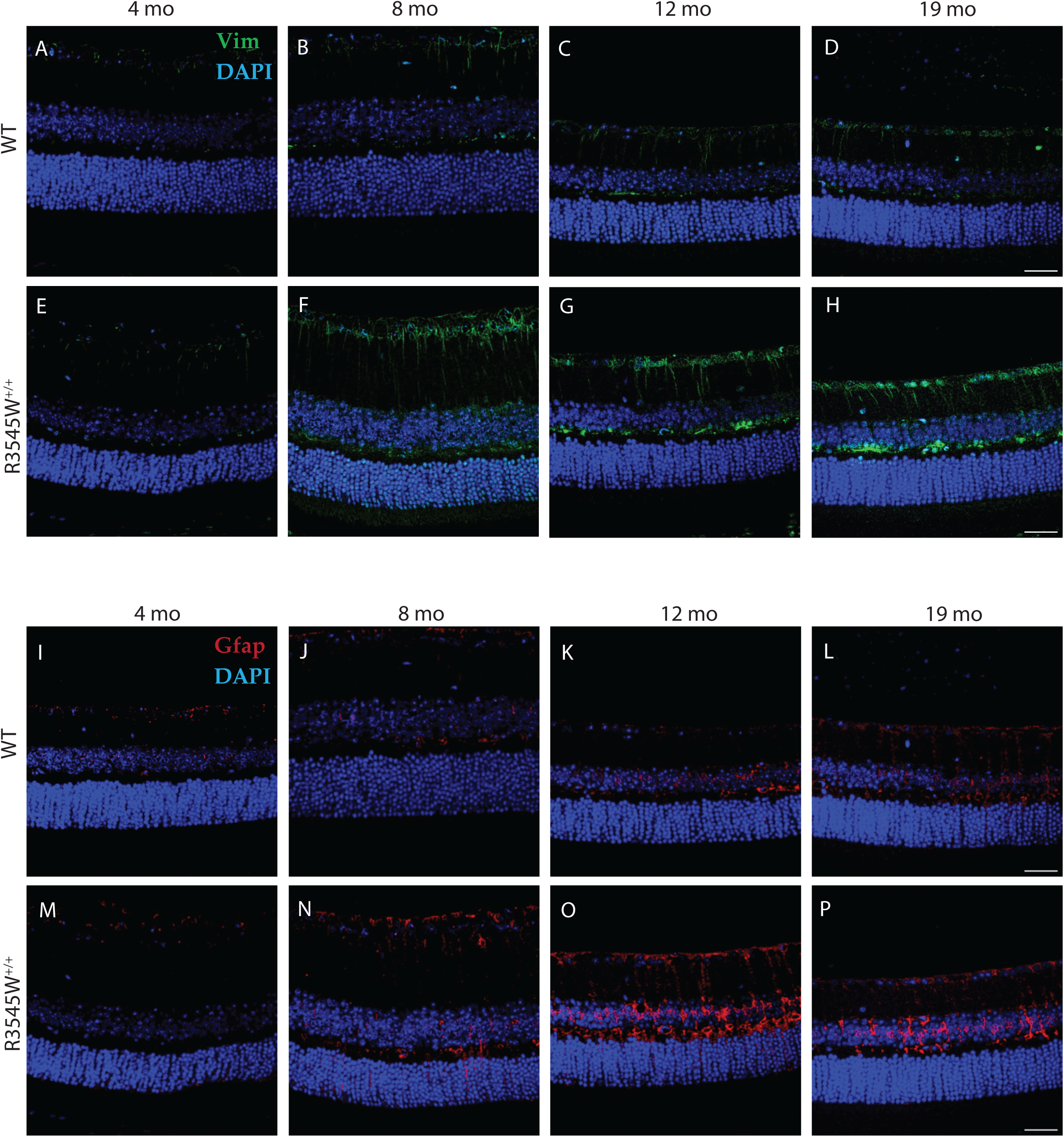
Muller glia driven gliosis is evident in aging R345W^+/+^ mice compared to WT. Representative images (n = 4-6 eyes) of retinal cross sections showing expression of vimentin (green) and GFAP (red) along with nuclear stain DAPI (blue) (scale bar = 50µm). Increased expression of vimentin and GFAP is observed in aging R345W^+/+^ mice, especially in the 8-12 mo window compared to their aging WT littermates.

Whereas Muller cells are the most common support glial cell in the eye, microglia are the resident immune cells of the retina and patrol various layers in response to cell damage or disease. In a young healthy state, Iba1^+^ microglial cells take on a ramified morphology and are located in the GCL, IPL, and OPL^96^. However, with disease or even age, Iba1^+^ cells can migrate from these layers (or extravagate from the circulation) into the subretinal space, including in the photoreceptor and RPE layers where they adopt a primed or ameboid-like morphology^97^. In aged WT mice (20 mo), Iba1^+^ cells appear primarily ramified in the GCL, IPL, and OPL (Fig. 6A-C), indicating a largely quiescent state. In contrast, Iba1^+^ cells in these same layers in R345W^+/+^ mice are hypertrophic with increased soma size (Fig. 6A-C), reflecting an activated state^98, 99^. Due to their advanced age, even WT mice had detectable infiltration of Iba1^+^ cells in the photoreceptor (PR) and RPE layers, which is not commonly found in young, healthy mice. However, in each of the cell layers quantified, R345W^+/+^ mice had significantly higher numbers of Iba1^+^ cells (Fig. 6A-E), indicating either increased proliferation of resident microglia or infiltrating monocytes from the circulation. Overall, the combination of these results suggest that the R345W mutation triggers a state of glial activation (both Muller glia and microglia) as well as subretinal homing of mobile Iba1^+^ immune cells.

**Fig. 6.**
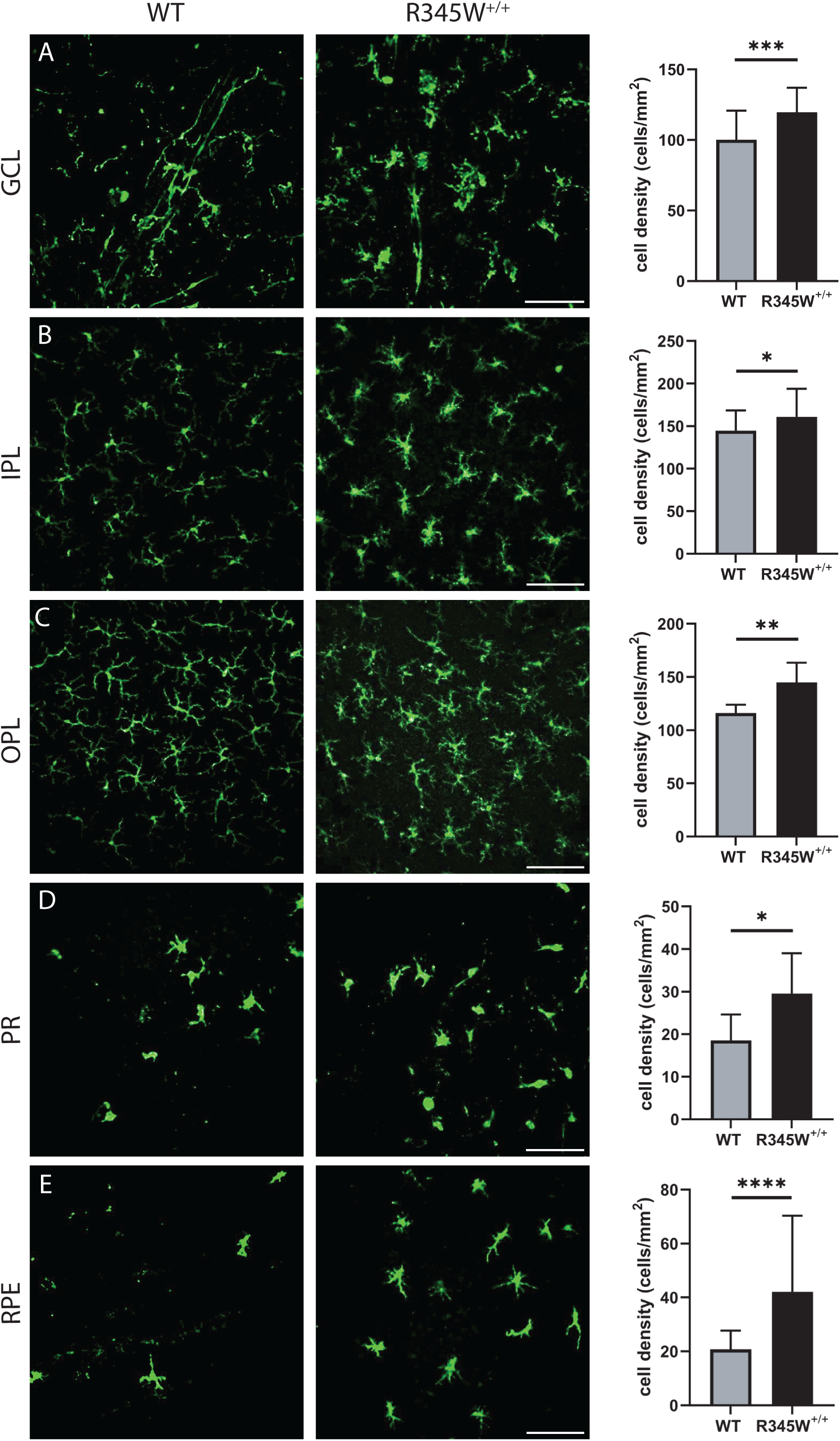
Microglia activation associated hypertrophy and proliferation observed in R345W^+/+^ mice compared to WT at 20 mo. (A-E) Representative images (n = 5-6 eyes) of Iba1 (green) stained microglia in the ganglion cell layer (GCL), inner plexiform layer (IPL), outer plexiform layer (OPL), photoreceptor layer (PR) and RPE (scale bar = 100 µm). Graphs representing cell densities in each of the specified layers (R345W^+/+^ vs WT) showing increased microglia in all layers quantified. Note: data presented here were analyzed using a Two-way ANOVA in combination with data presented in Fig. 9. * p < 0.05, ** p < 0.01, *** p < 0.001, **** p < 0.0001. Mean ± S.D. R345W^+/+^ images and quantification are identical to that shown later in Fig. 9.

### Knockout of Casp1 or Nlrp3 protects against R345W^+/+^-induced BLamD formation and growth

The NLRP3 inflammasome functions to regulate the innate immune responses by activating CASP1, which further cleaves pro-IL18 and pro-IL-1β inflammatory cytokines. Based on our transcriptional analysis (Fig. 3) and immunohistochemistry findings (Figs. 4, 6) in R345W^+/+^ mice, we hypothesized that mutant Efemp1 was triggering chronic sterile inflammation mediated through the Nlrp3 inflammasome and contributing to BLamD formation. Accordingly, we speculated that genetic elimination (or chemical inhibition) of the Nlrp3 pathway would reduce or prevent BLamD formation. To accomplish this work, we crossed R345W^+/+^ mice with Casp1^-/-81^ or Nlrp3^-/-100^ mice on the C57BL6 background^101, 102^. In addition to standard genotyping, as an initial step to verify functional removal of the Nlrp3 inflammasome, we isolated bone marrow-derived macrophages (BMDMs) from 2 mo mice followed by treatment with a canonical pathogen associated molecular pattern (PAMP, lipopolysaccharide) and/or damage associated molecular pattern (DAMP, nigericin) followed by an IL1β Lumit assay^103^ (Sup. Fig. 5A-D). Casp1^-/-^ R345W^+/+^ and Nlrp3^-/-^ R345W^+/+^ mice both demonstrated virtually no Nlrp3 inflammasome activity (Sup. Fig. 5C, D) when compared to WT or R345W^+/+^ BMDM (Sup. Fig. 5A, B).

After successfully verifying the functional inactivation of the Nlrp3 pathway in Casp1^-/-^ and Nlpr3^-/-^ BMDM, we sought to measure the impact that absence of this pathway had on BLamD initiation and propagation in the R345W^+/+^ genetic background. To perform these comparisons, we chose to focus on the 12 mo timepoint since this was the youngest age that provided the greatest difference between WT and R345W^+/+^ BLamD size (Fig. 2A-C). Excitingly, both Casp1^-/-^ R345W^+/+^ and Nlrp3^-/-^ R345W^+/+^ mice were significantly protected from the formation of BLamDs both with respect to size, but also coverage across the retina (Fig. 7A-D). In fact, the average BLamD size in Casp1^-/-^ R345W^+/+^ and Nlrp3^-/-^ R345W^+/+^ mice at 12 mo was not only significantly less than those found in R345W^+/+^ mice, but actually approached (or was lower than) WT BLamD levels (Fig. 7A-D, mean BLamD size in WT mice = 4.39 μm^2^, R345W+/+ = 15.88 μm^2^, Casp1^-/-^ R345W^+/+^ = 5.97 μm^2^, and Nlrp3^-/-^ R345W^+/+^ = 2.86 μm^2^). These results suggest that the NLRP3 inflammasome and its downstream effectors are key contributors to the pathological formation of BLamD in R345W^+/+^ mice, and possibly ML/DHRD more broadly.

**Fig 7.**
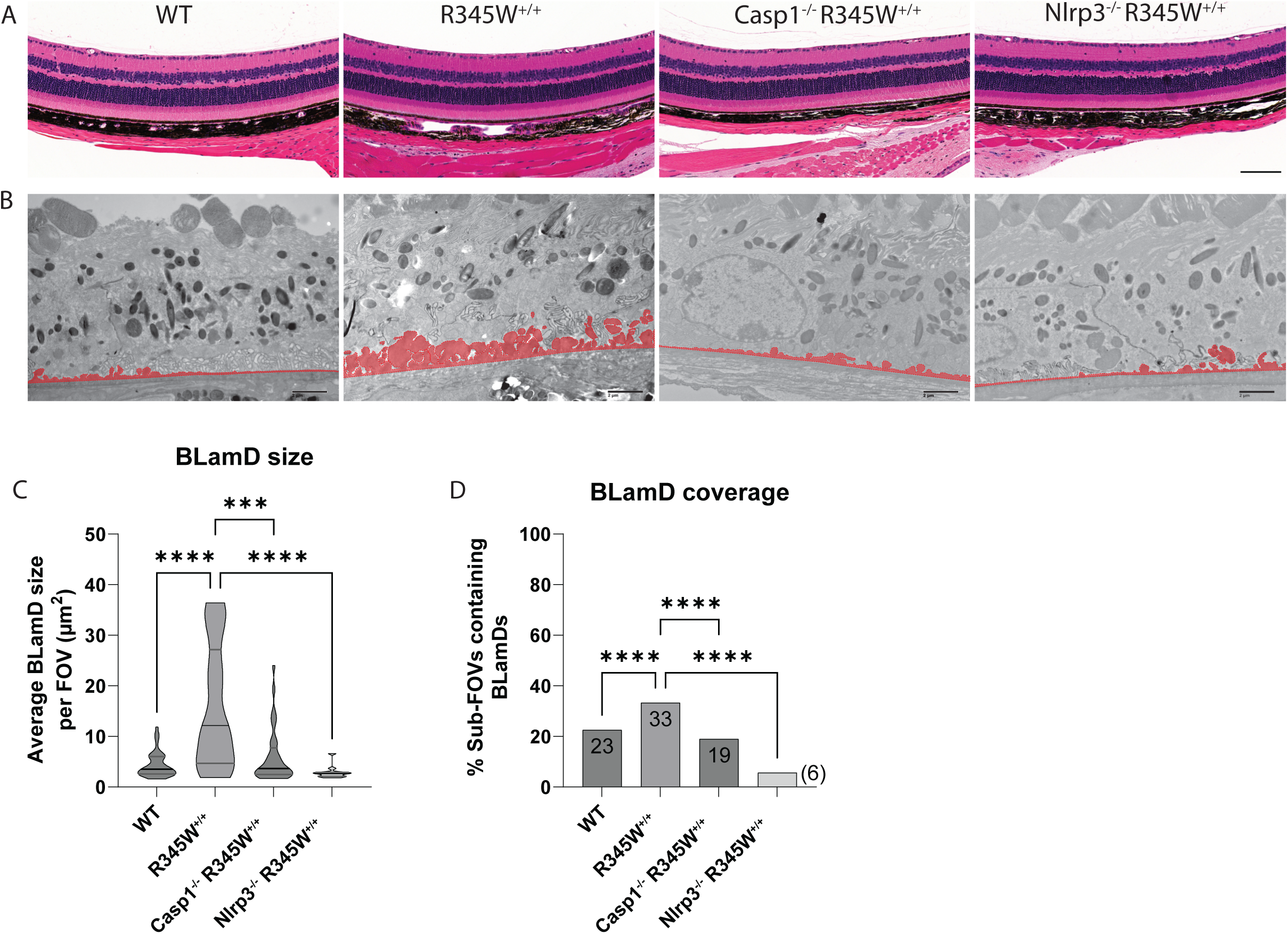
Retinal histology and BLamD comparison of WT, R345W^+/+^, Casp1^-/-^ R345W^+/+^, and Nlrp3^-/-^ R345W^+/+^ 12 mo mice. (A) H&E-stained retinal histology sections demonstrates that removal of Casp1 and Nlrp3 is well tolerated structurally. Scale bar = 100 µm. (B) Representative BLamD traced electron micrographs from each genotype. Scale bar = 2 µm. (C) Violin plot depicting comparative analysis of BLamD size per FOV across different genotypes. (D) Comparative analysis of the abundance of sub-FOVs containing BLamDs presented as percent coverage across BrM. n = 3 males/3 females for each genotype. Analysis performed using Kruskal-Wallis test. Note: TEM images of WT and R345W^+/+^ mice as well as quantification (C, D) are identical to the 12 mo age group displayed earlier in Fig. 2.

### Nlrp3 deficiency reduces microglia infiltration/proliferation in the R345W^+/+^ background and broadly mitigates increases in cytokines/chemokines observed in the RPE and retina, but does not affect Muller cell gliosis

Given the pronounced effects of removing Nlrp3 on BLamD size and retinal coverage in the R345W^+/+^ background, we decided to focus on this knockout strain moving forward instead of pursuing both Casp1^-/-^ R345W^+/+^ and Nlpr3^-/-^ R345W^+/+^ mice simultaneously. To gain a better understanding of how seemingly pathological features observed in the R345W^+/+^ mice (i.e., Muller cell gliosis, Iba1^+^ activation/infiltration/proliferation, and generalized inflammation) correlate with or influence BLamD formation and are potentially altered in Nlrp3^-/-^ mice, we performed a series of immunohistochemistry and molecular analyses in Nlrp3^-/-^ and Nlrp3^-/-^ R345W^+/+^ mice. First, we assessed Vim and Gfap staining intensity in 12 mo Nlrp3^-/-^ and Nlrp3^-/-^ R345W^+/+^ mice (Fig. 8A-D). There was minimal but detectable Gfap and Vim staining in Nlrp3^-/-^ mice, with staining patterns and intensities similar to those observed in WT mice at this age (Fig. 5C, K). Surprisingly, Nlpr3^-/-^ R345W^+/+^ mice still demonstrated high levels of both Gfap and Vim, bearing staining patterns that resembled R345W+/+ mice at this age (Fig. 5G, O).

**Fig. 8.**
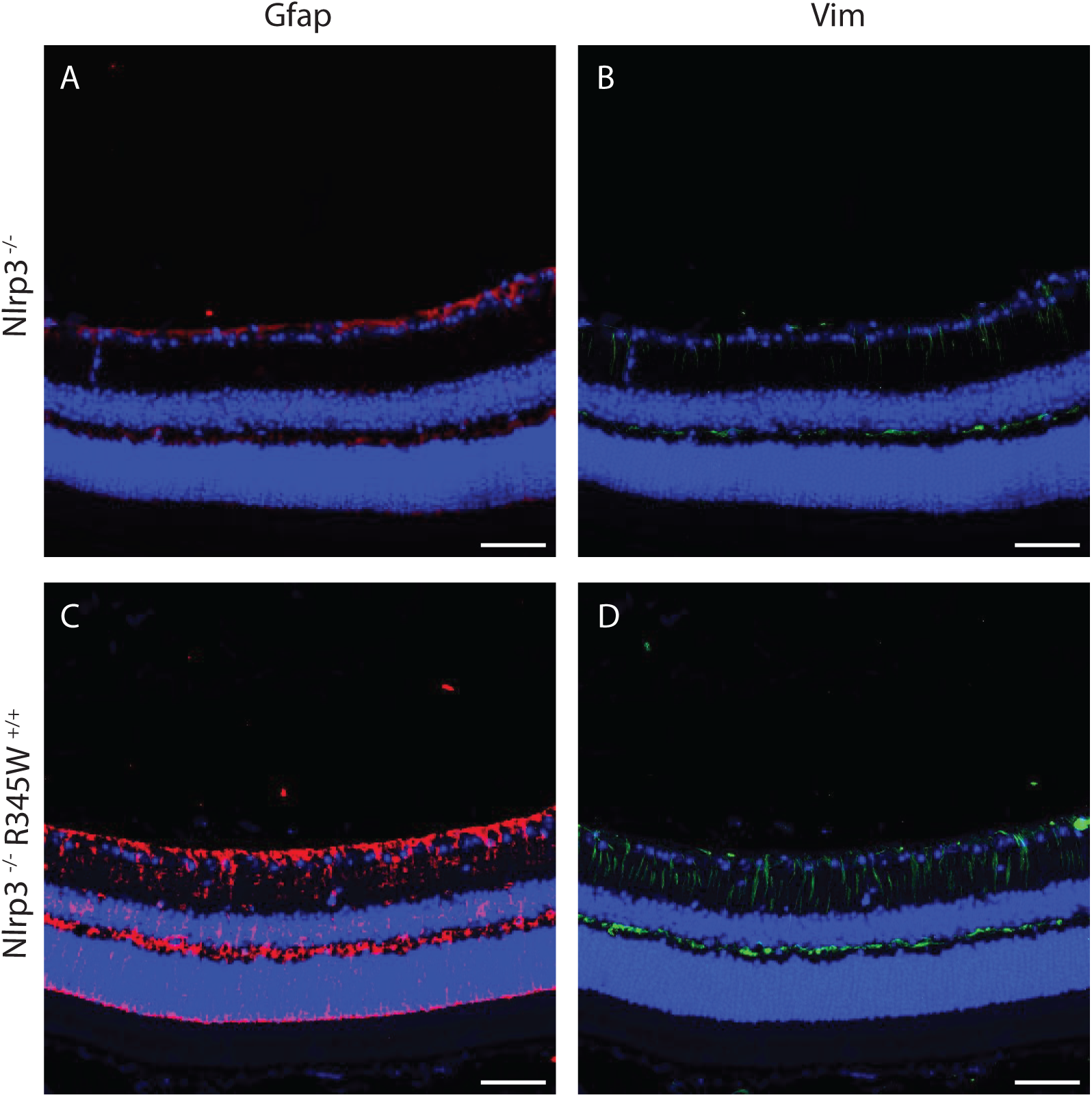
Muller cell gliosis is unaffected by the absence of Nlrp3 in R345W^+/+^ mice at 12 mo. (A-D) Representative images (n = 4) of retinal cross sections showing expression of Gfap (A, C, red) and Vim (B, D, green) along with nuclear stain DAPI (blue) as a reference (scale bar = 50 µm). Increased expression of Gfap and Vim is observed in Nlrp3^-/-^ R345W^+/+^ mice (similar to that seen in R345W+/+ mice, cf. Fig. 5G, O) compared to Nlrp3^-/-^ (or WT, cf. Fig. 5C, K) mice.

Although R345W-triggered Muller cell gliosis appeared to be operating through Nlrp3 independent pathways, genetic removal of Nlrp3 significantly affected Iba1^+^ cell density and morphology throughout the retina (Fig. 9A-E). Within the GCL, IPL, and OPL, Iba1^+^ microglia appeared to revert to a ramified, non-activated state (Fig. 9A-C) similar to what was observed in WT mice (see Sup. Fig. 6A-E for full comparison of WT, R345W^+/+^ and Nlrp3^-/-^ R345W^+/+^ Iba1 staining). Iba1^+^ microglia soma were noticeably smaller in Nrlp3 deficient mice (Fig. 9A-C) and there was a significant reduction in subretinal infiltrating microglia in both the PR and the RPE cell layers (Fig. 9D, E). To complement these immunostaining studies, we also performed a discovery-based cytokine/chemokine/growth factor array assay in WT, R345W^+/+^ and Nlrp3^-/-^ R345W^+/+^ RPE and NR tissue homogenates (Sup. Fig. 7A-D). We found a series of proteins that were upregulated in R345W^+/+^ tissue (Sup. Fig 7A, B) which were ‘normalized’ to WT levels through elimination of Nlrp3 (Sup. Fig. 7C, D), which generally supported the notion of a proinflammatory retinal environment in R345W^+/+^ mice that is resolved or minimized by removal of Nlrp3. Specific cytokines that were upregulated in both the RPE and NR in R345W^+/+^ mice and subsequently downregulated by removal of Nlrp3 include immune cell modulators proliferin^104^, Reg3G^105^, and WISP-1^106^ (Sup. Fig. 7A-D) which corroborates our observations of reduced Iba1^+^ microglia activation and recruitment/proliferation in Nlrp3^-/-^ R345W^+/+^ mice (Fig. 9A-E).

**Fig. 9.**
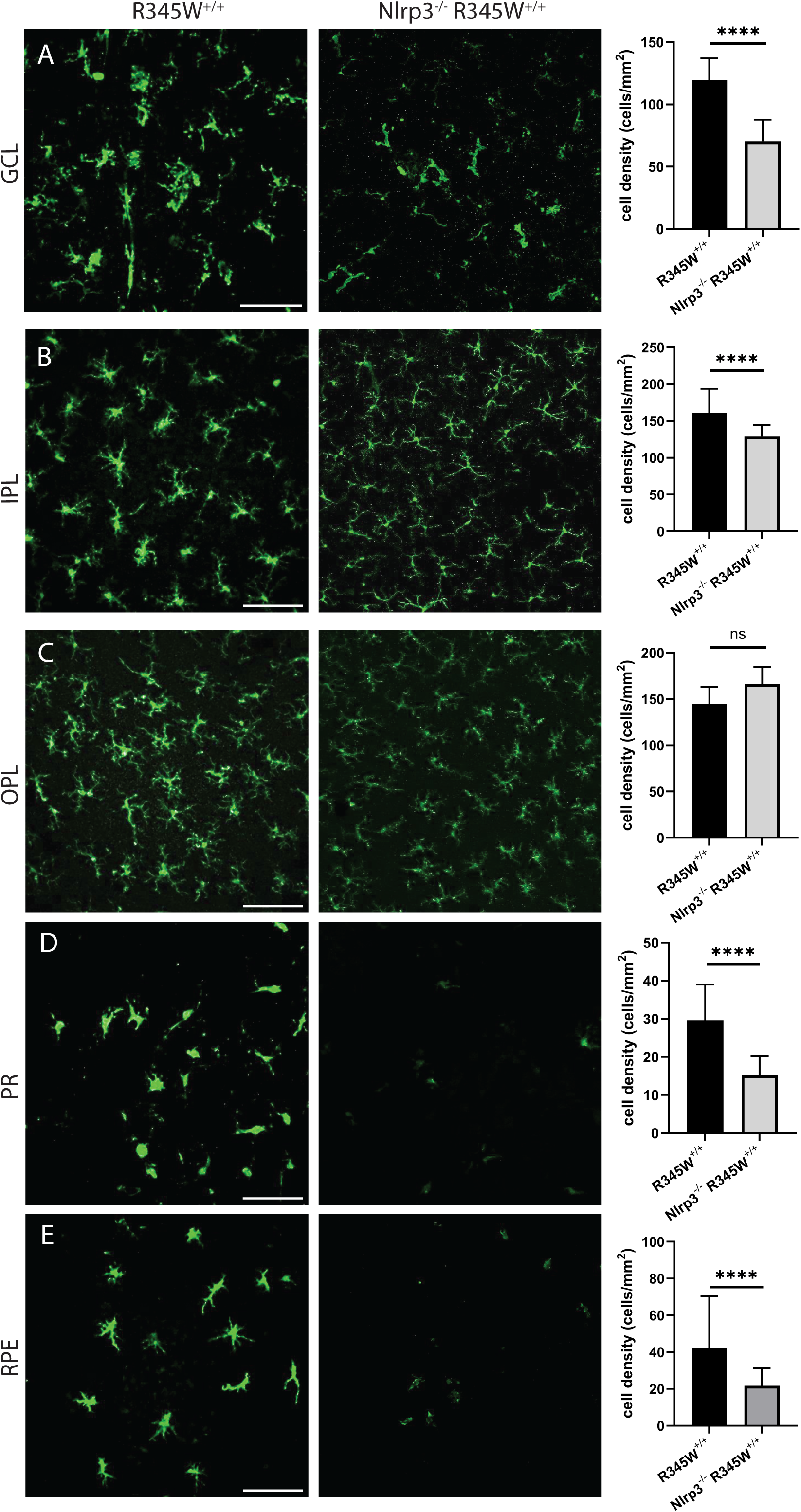
Microglia activation-associated hypertrophy and proliferation due to the R345W mutation is prevented in Nlrp3 deficient mice. Nlrp3^-/-^ R345W^+/+^ mice. (22 mo) mice compared to R345W^+/+^ (20 mo). Representative images (n = 4-6) of Iba1^+^ (green) stained microglia in the ganglion cell layer (GCL), inner plexiform layer (IPL), outer plexiform layer (OPL), photoreceptor layer (PR) and RPE (scale bar = 100 µm). Graphs representing cell densities in each of the specified layers showing significant reductions in microglia densities in the GCL, IPL, PR, and RPE layers. Note: data presented here were analyzed using a two-way ANOVA in combination with data presented in Fig. 6. **** p < 0.0001. Mean ± S.D. Note: R345W^+/+^ images and quantification are identical to that shown earlier in Fig. 6.

### Genetic elimination of Nlrp3 prevents age-related spontaneous BLamD formation in WT mice

A major motivating factor of studying the R345W ML/DHRD retinal dystrophy mouse model is the hope to leverage findings in this rare retinal dystrophy for understanding and treatment of idiopathic early/intermediate AMD, which affects nearly 180 million people worldwide^29^. Thus, based on our observation that Nlrp3 deficiency strongly protects against the R345W-induced BLamD phenotype, we next asked whether Nlrp3 removal could also protect against spontaneous, idiopathic BLamD formation in WT mice. Twelve-month-old WT, Casp1^-/-^, or Nlrp3^-/-^ mice all demonstrated healthy retinas as assessed by histology (Fig. 10A). As we previously demonstrated, WT mice at this age do form sporadic BLamDs that continue to grow during aging (Fig. 2, Sup. Fig. 4). Removal of Casp1 led to slightly lower, but not statistically different BLamD size and coverage (Fig. 10B-D). However, excitingly, Nlrp3 removal led to significant protection against BLamD size and coverage in the WT background (Fig. 10B-D), reaffirming the idea that cellular pathways identified and validated as protective in the R345W ML/DHRD mouse have the promise of also protecting in more prevalent diseases such as idiopathic AMD.

**Fig. 10.**
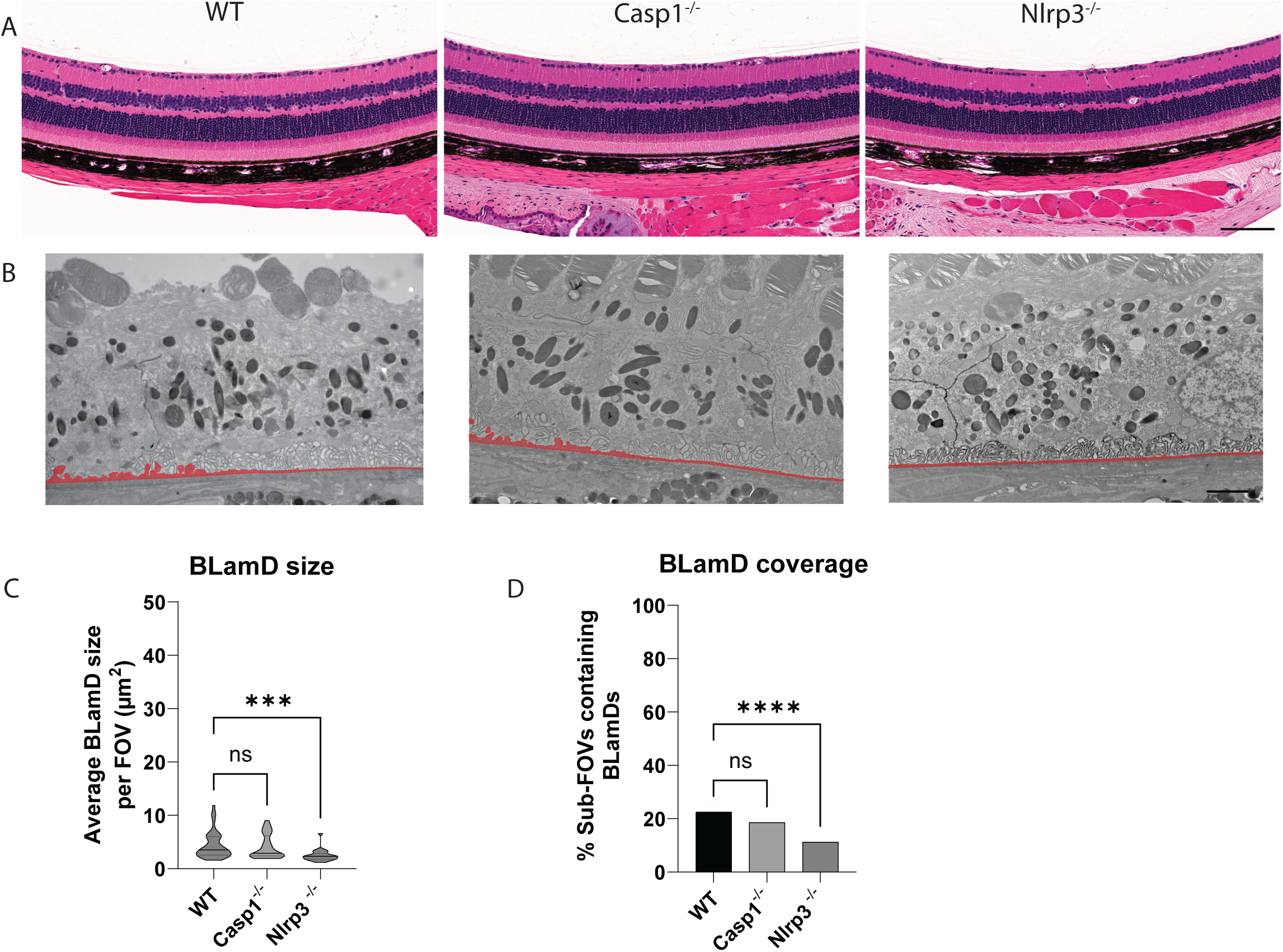
Nlrp3 deficiency protects mice from spontaneous BLamD deposit formation. (A) 12 mo H&E-stained histology images of WT, Casp1^-/-^, and Nlrp3^-/-^ mice. Scale bar = 100 µm. (B) Representative TEM images from 12 mo mice. (C, D) quantitation of BLamD size (µm^2^/FOV) and percent sub-FOVs containing BLamDs. Scale bar = 2 μm Comparative analysis performed using Kruskal-Wallis test. **** p < 0.0001. Note: The WT TEM image and WT BLamD quantification are identical to that shown previously in Fig. 2A-E.

## DISCUSSION

ML/DHRD is a clinically heterogenous macular dystrophy with surprisingly variable intra- and interfamilial variability with respect to human disease severity and onset^74, 107^. These results suggest that ML/DHRD disease trajectory or initiation is likely strongly affected by environmental cues or subtle genetic differences among individuals. Accordingly, small molecules or genetic perturbations that alter key cellular responses involved in ML/DHRD progression (e.g., EFEMP1 protein production/turnover, or stress responses due to malformed EFEMP1) have the promise of serving as first-in-human therapies for this rare disease which may comprise up to 0.9% of all inherited retinal dystrophies (IRDs) in select populations^108^, amounting to thousands of individuals worldwide.

Our work clearly defines an important role for the Nlrp3 inflammasome in mediating BLamD pathology in the R345W mouse model of ML/DHRD and in aging wild-type mice. These results add to previous findings that Nlrp3/NLRP3 mediates RPE atrophy in mouse models of AMD and in human AMD donor tissue^19, 38, 39^. In further support of inflammasome involvement in the pathology of AMD, recently a 38 yo Chinese female with a constitutively active, autosomal dominant autoimmune mutation in NLRP3 (c.1043C>T, p.T348M) presented with pseudodrusen (amorphous material between the RPE and photoreceptors) and hard drusen near the optic nerve^46^. Accordingly, based on the culmination of these data, it is tempting to speculate that compounds such as MCC950^109^, a potent nanomolar direct NLRP3 inhibitor, may serve as promising leads for ML/DHRD therapeutics and possibly sporadic AMD. Yet, while many NLRP3 inhibitors are effective systemically on circulating immune cells, they suffer from poor penetration into the brain or eye^110^ and have been found to cause liver toxicity^111, 112^. Future studies focused on developing a better understanding regarding the BLamD-related contributions of ocular NLRP3 vs. systemic NLRP3 will provide more insight on whether NLRP3 needs to be inactivated directly in the eye (and where in the eye^113^), in the circulation, or both.

Canonical triggering of innate immunity via full NLRP3 activation follows a two-step mechanism involving a ‘priming’ step typically with a pathogen-derived PAMP followed by an ‘activation’ step with a damage- or danger-associated DAMP. In the instance of the ML/DHRD mouse, it is unlikely that a canonical PAMP (i.e., cellular components from bacteria/protozoa/fungi or viral DNA/RNA) is present in the eye, begging the question of whether a PAMP is indeed required for ocular NLRP3 triggering. Alternatively, it is possible that NLRP3 is triggered during sterile (non-infectious) inflammation, as suggested/demonstrated previously^114, 115, 116^. Based on what is known about the molecular consequences of R345W expression, there are a series of potential DAMPs that could serve as a source for activating NLRP3. For example, we and others have demonstrated that R345W mice upregulate complement component 3 (C3)^75^, which is ultimately cleaved to C3a. This cleavage product in turn can increase ATP efflux and NLRP3 activation^117^. Notably, genetic elimination of C3 prevented BLamD formation in R345W mice^75^ while removal of C5 had no effect^76^. Based on these findings, it is interesting to speculate that perhaps FDA-approved C3 inhibitors (for geographic atrophy) such as Pegcetacoplan (SYFOVRE)^118^ may ultimately be repurposed for use in ML/DHRD, possibly with better functional outcomes than has occurred in dry AMD. Another potential source of inflammasome activation in the ML/DHRD mouse could originate from cholesterol crystals^119^. The R345W mutant has been demonstrated by some groups to impair cholesterol efflux from the RPE, leading to higher intracellular esterified cholesterol levels^58^. Each of these pathways may serve as therapeutic targets for ML/DHRD in the near future.

Aside from identifying a new and small molecule-targetable pathway for minimizing ML/DHRD pathology, our work also provides a quantitative time course of BLamD initiation and growth in WT and R345W^+/+^ mice from 4 mo to 20 mo. These data are important from at least two prospectives. First, they provide an actionable window during which therapeutics can be administered with greatest effect. For example, between 8 mo and 12 mo, R345W^+/+^ BLamD size spike and subsequently plateau. Treating mice at 8 mo of age, as we have performed previously with a GSK3 inhibitor^57^, before this large spike will likely enable better evaluation of a therapeutic effect. While this same window is not as well defined in WT mice, there is a substantial increase in BLamD coverage between 16 and 20 mo that would likely afford the best dynamic range for evaluating potential therapies. Along these lines, defining biologically what is occurring in the retina/RPE to initiate these dramatic effects is of significant interest. Second, we believe that our quantitative approach provides a more accurate representation of BLamD formation than the representative images normally provided in studies. Based on previous representative images^67, 75^, it would appear that WT mice don’t form any appreciable BLamDs, even with advanced age. However, when approaching this topic from a more systemic and quantitative manner, we were surprised to find that indeed they do form BLamDs, and by 20 mo, the coverage of BLamDs is not different than what is found in R345W^+/+^ mice, albeit smaller. These findings allowed us to use WT mice to evaluate whether Nlrp3 deficiency also prevented sporadic BLamD formation - results that bear promise potentially for sporadic AMD.

Yet the cellular contributions that influence BLamD formation remain unclear. Our data suggest that expression of R345W Efemp1 (which is produced in Muller glia cells according to Spectacle^120^, a single cell RNA sequencing visualization database) induces Muller cell gliosis starting at 8 mo and peaking at 12 mo. While removal of Nlrp3 had no effect on R345W-induced Muller cell gliosis, it did significantly protect against BLamD formation at this age. These results suggest that Muller cell gliosis in this instance is likely responding to an altered ECM produced by the R345W mutant and does not correlate with (and may not contribute to) BLamD formation in mice. However, Iba1^+^ microglia infiltration into the subretinal (PR and RPE) space did correlate with BLamD formation – the more Iba1^+^ activated microglia in the subretinal space paralleled BLamD formation. Nlrp3 removal significantly reduced Iba1^+^ cells in the subretinal space and protected against BLamD formation in WT and in R345W^+/+^ mice. Future studies will begin to unravel whether the infiltrating Iba1^+^ microglia are resident cells that have proliferated and migrated to the subretinal space or infiltrating monocytes from the circulation which have been differentiated into microglia. Moreover, we will begin to determine whether microglia have a more direct effect on production of BLamD formation due to their ability to produce and remodel the local ECM^121, 122^, or whether the presence of BLamD drive the recruitment of microglia in R345W^+/+^ and aging WT mice.

Overall, our findings on the importance of Nlrp3 in BLamD formation adds to the growing body of research identifying potential therapeutic routes to alleviate or delay pathology in ML/DHRD^57, 58, 75^, and support the idea that select retinal dystrophies with phenotypic similarities to AMD can be likely used as proving grounds to develop breakthroughs in etiologically complex, yet prevalent diseases such as sporadic AMD.

## ACKNOWLEDGEMENTS

We thank Randy Nessler, Director of the University of Iowa Central Microscopy Research Facility for training AJO on how to process mouse ocular tissue for transmission electron microscopy analysis. JDH is the Larson Endowed Chair for Macular Degeneration Research (UMN) and is supported by the Helen Lindsay Foundation, the Edward N. and Della L. Thome Memorial Foundation Award in Age-Related Macular Degeneration Research, and R01-EY027785. HZ is supported by R01-DK125352 and R01-DK128031. Work described herein was also supported in part by the UT Southwestern Peter O’Donnell Jr. Brain Institute (OBI) Sprouts Pilot Grant (SD).

## SUPPLEMENTAL FIGURE LEGENDS

<u>**Sup. Fig. 1.** RPE subpopulation morphometry is similar in WT and R345W^+/+^ mice.</u> (A-D) Graphs comparing cell area (A), aspect ratio (B), hexagonality (C), and neighbors (D) for the three RPE subpopulations in WT and R345W^+/+^ at 22 mo. To assess the statistical significance of the observed differences in morphometric features among the subpopulations, a two-way ANOVA was performed. Mean ± S.D., n = 6 for WT, n = 7 for R345W^+/+^.

<u>**Sup. Fig. 2.** Corneal opacity, IOP, and RGC density are similar between WT and R345W^+/+^ mice.</u> (A) Unremarkable representative slit lamp images from WT and R345W^+/+^ mice (14 mo, n = 5-6 mice). (B) IOP measurements in WT and R345W^+/+^ mice (14 mo, n = 5-6 mice; 10-12 eyes) are not statistically different. Unpaired t-test. Mean ± S.D. (C) RGC density in WT and R345W^+/+^ as determined by RBPMS staining (17 mo, n = 7-8 eyes) is also not different between genotypes. DAPI (blue) was used as nuclear stain (Scale bar = 100 µm). (D) Graph representing RGC cell densities that remain unchanged in R345W^+/+^ mice compared to WT. Unpaired t-test. Mean ± S.D.

<u>**Sup. Fig. 3.** Serum cholesterol, triglycerides, and body composition are not different in WT and mutant F3 mice.</u> (A) cholesterol or (B) triglyceride analysis was performed from serum obtained from 16 mo mice, n = 2-5, mean ± S.E.M. (C) Body composition was evaluated by a micro-CT scanner (Bruker) at 12 mo (n = 6-10 individual mice). Only statistical differences in body composition were observed in the mass of male vs. female mice. One-way ANOVA. * p < 0.05, ** p < 0.01.

<u>**Sup. Fig. 4.** Re-presentation of data from Fig. 2 by genotype and BLamD size/coverage.</u> (A-D) to obtain a better longitudinal view of BLamD size and coverage (growth), data from Fig. 2 was re-plotted according to genotype and across ages. Two-way ANOVA with Kruskal-Wallis test for multiple comparisons. **** p < 0.0001. n = 6 mice/genotype and age group (3 male, 3 female) except for 20 mo mice (n = 3 females/genotype). Median along with 1^st^ and 3^rd^ quartiles plotted.

<u>**Sup. Fig. 5.** Confirmation of functional impairment of the Nlrp3 inflammasome using BMDM.</u> (A-D) An IL1β Lumit assay was used to evaluate cytokine production in differentiated BMDM isolated from 2 mo mice after treatment with LPS and/or nigericin. 8 mice in total used, all female. One mouse for each genotype. 6 replicate wells for each treatment group and the experiment was repeated twice and data combined.

<u>**Sup. Fig. 6.** Re-presentation of data from Fig. 6 and Fig. 9 in totality.</u> Since data from these two figures were analyzed together, we present the data from Fig. 6 and from Fig. 9 together for comparison. Two-way ANOVA. * p < 0.05, ** p < 0.01, *** p < 0.001, **** p < 0.0001. Mean ± S.D.

<u>**Sup. Fig. 7.** Cytokine array analysis of RPE/choroid and neural retinal (NR) tissues in WT, R345W^+/+^, and Nlrp3^-/-^ R345W^+/+^ mice.</u> (A-D) Samples were collected from perfused 12 mo WT, R345W^+/+^, and Nlrp3^-/-^ R345W^+/+^ female mice. In total, 111 cytokines were assayed in duplicate for each tissue sample from all genotypes. The top 11 cytokines in the RPE/choroid (A) and top 8 cytokines measured in the NR (B) with the greatest fold change in expression were highlighted. These same cytokines were all downregulated in RPE/choroid (C) and NR (D) in Nlrp3^-/-^ R345W^+/+^ samples, reverting their abundance to near WT-levels in many cases. Cytokines spotted in duplicates. One mouse per genotype. 8 mice total used for these arrays.

## SUPPLEMENTAL MATERIALS AND METHODS

### Conscious intraocular pressure (IOP) measurement

Mice were gently restrained by first placing them in a soft, clear plastic cone (DecapiCone, Braintree Scientific, Braintree, MA) and then securing them in a custom-made restrainer. This restrainer was positioned on a height-adjustable platform. After allowing a few minutes for acclimation, conscious IOPs were measured during the daytime using a TonoLab rebound tonometer (Colonial Medical Supply, Franconia, NH).

### Slit-lamp

Slit-lamp imaging was performed on anesthetized mice (2.5% isoflurane and 0.8 L/min oxygen). Anterior chamber phenotypes were evaluated using a slit lamp (Haag-Streit Diagnostics, Mason, OH) and recorded with a digital camera (Canon EOS Rebel T6 DSLR, 18.0-megapixel CMOS image sensor and DIGIC 4+ Image Processor). All images were captured using consistent camera settings for brightness and contrast, with the "auto-focus" option enabled.

### Body composition analysis

Differences in mice body composition were assessed using a Bruker MiniSpec mq20 (Bruker Optics, Billerica, MA). Compositional changes in total body weight, body fat weight, lean tissue weight, fluid weight, percent weight of fat, and percent weight of lean tissue were assessed. Mice were scanned twice per session and the values were then averaged and subjected to comparison across genotypes using a two-way ANOVA and Tukey’s multiple comparison test. n = > 3.

### Bone marrow-derived macrophage (BMDM) isolation and treatment

To validate Nlrp3 inflammasome pathway inactivation (i.e., in mice lacking Nlrp3 or Casp1), we tested the release of the proinflammatory cytokine, IL-1β from BMDM cells originating from mice of each genotype. Femoral and tibial bone marrow from 2-month-old mice from each cohort were collected, cultured, and differentiated for 6 d into BMDMs using Iscove’s Modified Dulbecco’s Medium (IMDM) medium (ThermoFisher, Waltham, MA) supplemented with 10% FBS (Omega Scientific, Tarzana, CA), 25 ng/mL macrophage colony stimulating factor (M-CSF, ThermoFisher), 1% nonessential amino acids (ThermoFisher), and 1% penicillin-streptomycin (ThermoFisher). Cells were then replated onto a 12 well plate at 1.2 x 10^6^ cells/well and cultured overnight (Corning, Corning, NY). Cells were then subject to one of four treatments, 3 h 500 ng/µL lipopolysaccharide (LPS, Sigma, St. Louis, MO), 1 h 10 µM nigericin (Cayman, Ann Arbor, MI), a combination of LPS and nigericin, or untreated. IL-1β release into the cell culture media was assayed using a Lumit Mouse IL-1β Immunoassay (Promega, Madison, WI) followed by reading on a GloMax plate reader (Promega).

### Cytokine Profiling Array

To gain further insight into inflammation-related pathways potentially involved in BLamD formation, we took an unbiased cytokine/chemokine profiling approach. Twelve-month-old WT, R345W^+/+^, Casp1^-/-^ R345W^+/+^, Nlrp3^-/-^ R345W^+/+^ mice were deeply anesthetized using an i.p. ketamine/xylazine injection. Once unresponsive to foot pinch, the mice were perfused with 20 mL of heparinized saline to fully clear blood from the choroid. Eyes were immediately enucleated and dissected by cutting along the ora serrata to remove the anterior portion of the eye and the lens. The neural retina (NR) and retinal pigmented epithelium (RPE)/eye cups were then carefully separated, homogenized in PBS with protease inhibitors (ThermoFisherScientific) and 1% triton X-100 (Sigma-Aldrich) and frozen overnight at -80°C. Thawed samples were centrifuged and total sample protein was normalized across samples to 100 µg for NR and 200 µg for RPE/eyecups. Samples were then assayed using the Mouse XL Cytokine Array Kit ARY028 (R&D Systems, Minneapolis, MN) and imaged by following the Proteome Profiler Arrays with LI-COR Detection protocol (R&D Systems). The cytokine levels were quantified using the Protein Array Analyzer for ImageJ add-on by Gilles Carpentier (http://image.bio.methods.free.fr/ImageJ/?Protein-Array-Analyzer-for-ImageJ).

