## Supplementary figures and images for "Genetic removal of Nlrp3 protects against sporadic and R345W Efemp1-induced basal laminar deposit formation"

### Supplemental Fig 1

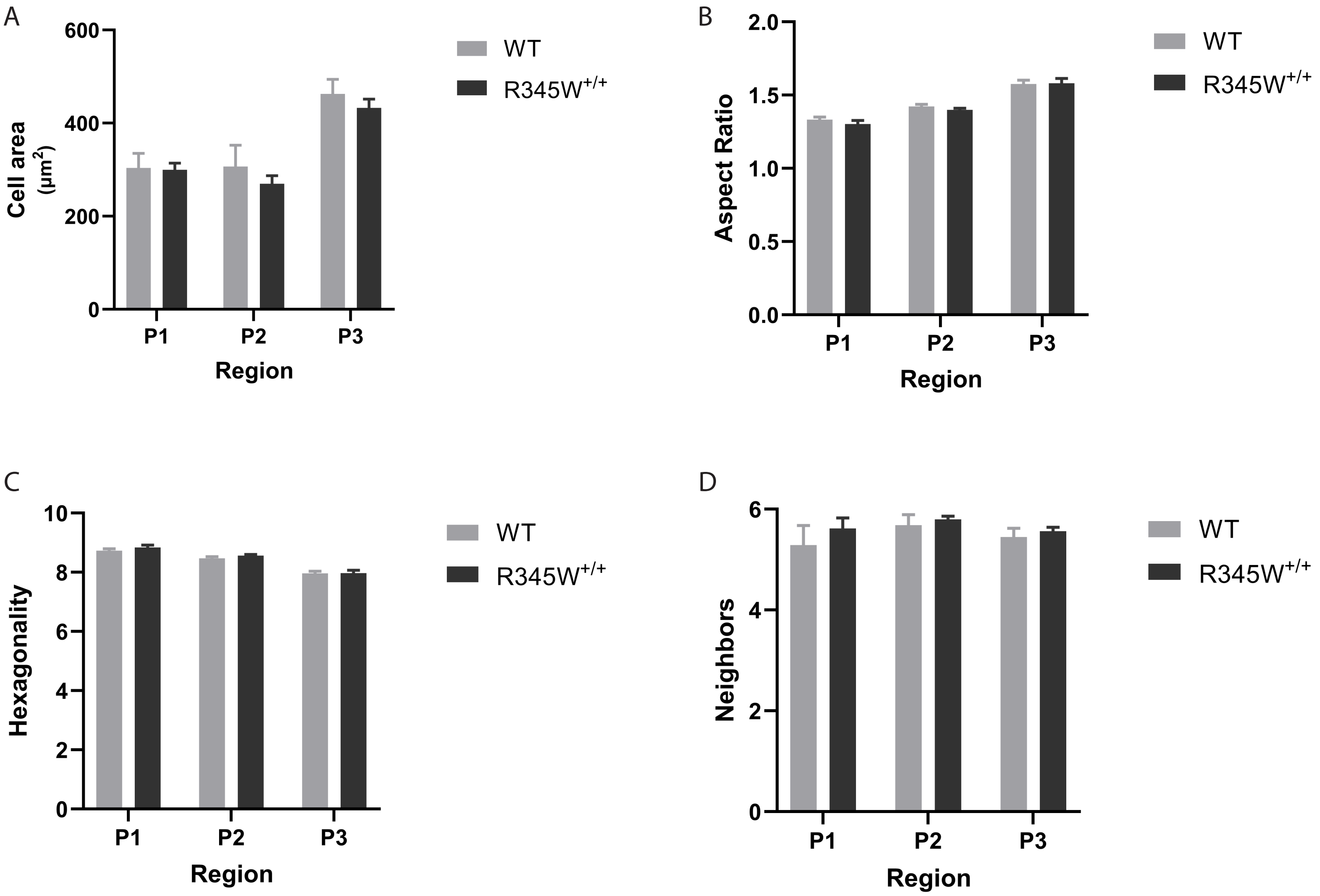

### Supplemental Fig 2

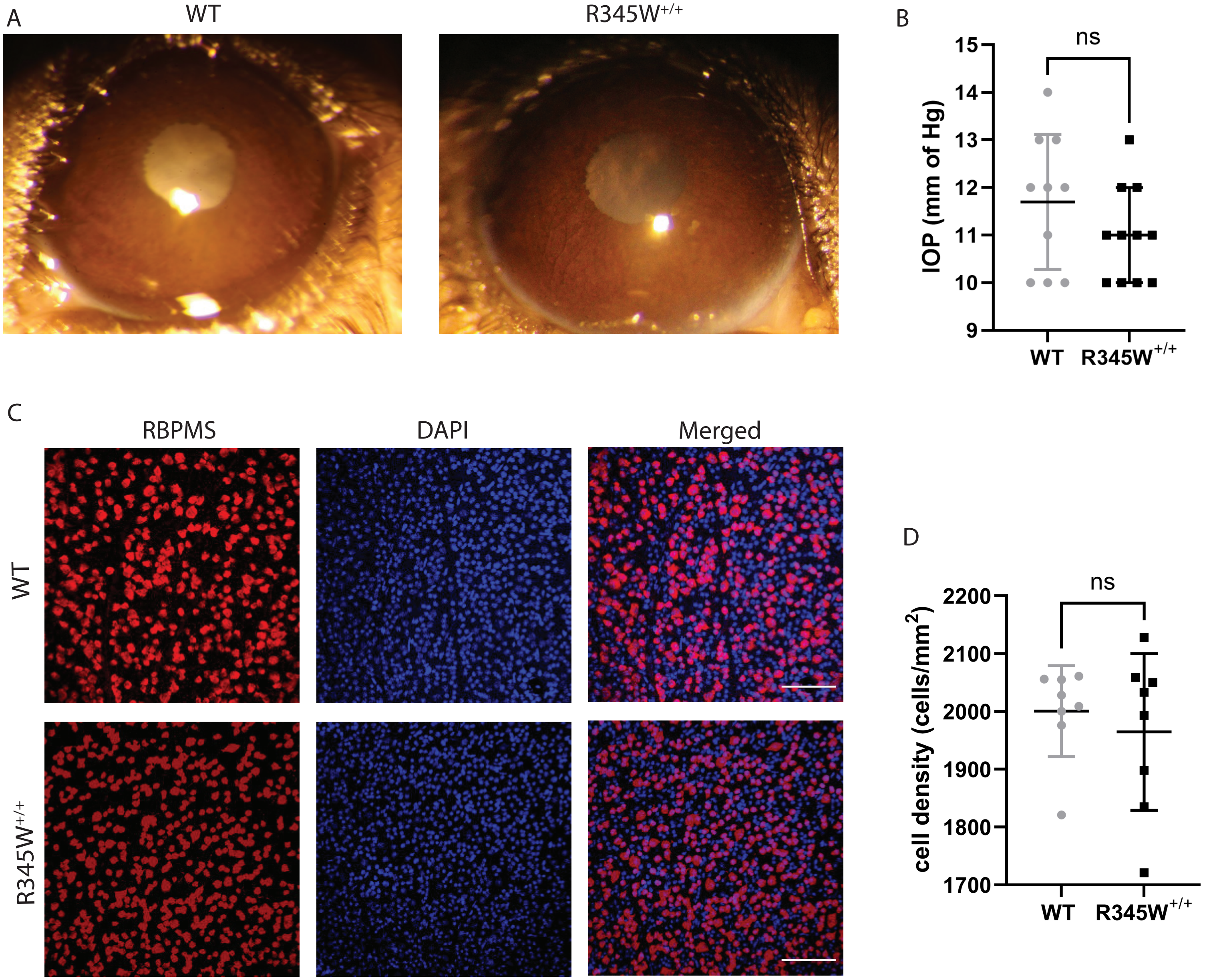

### Supplemental Fig 3

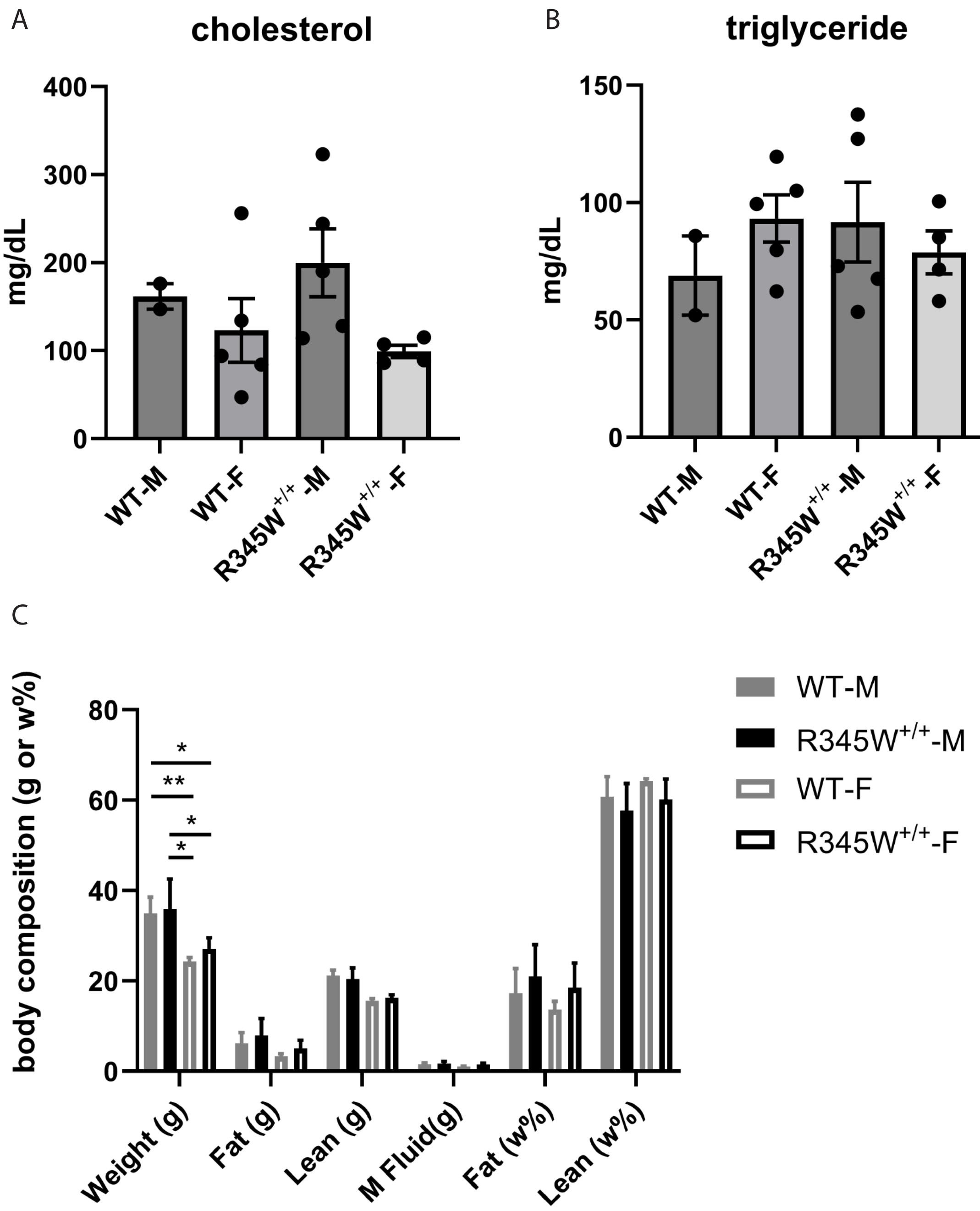

### Supplemental Fig 4

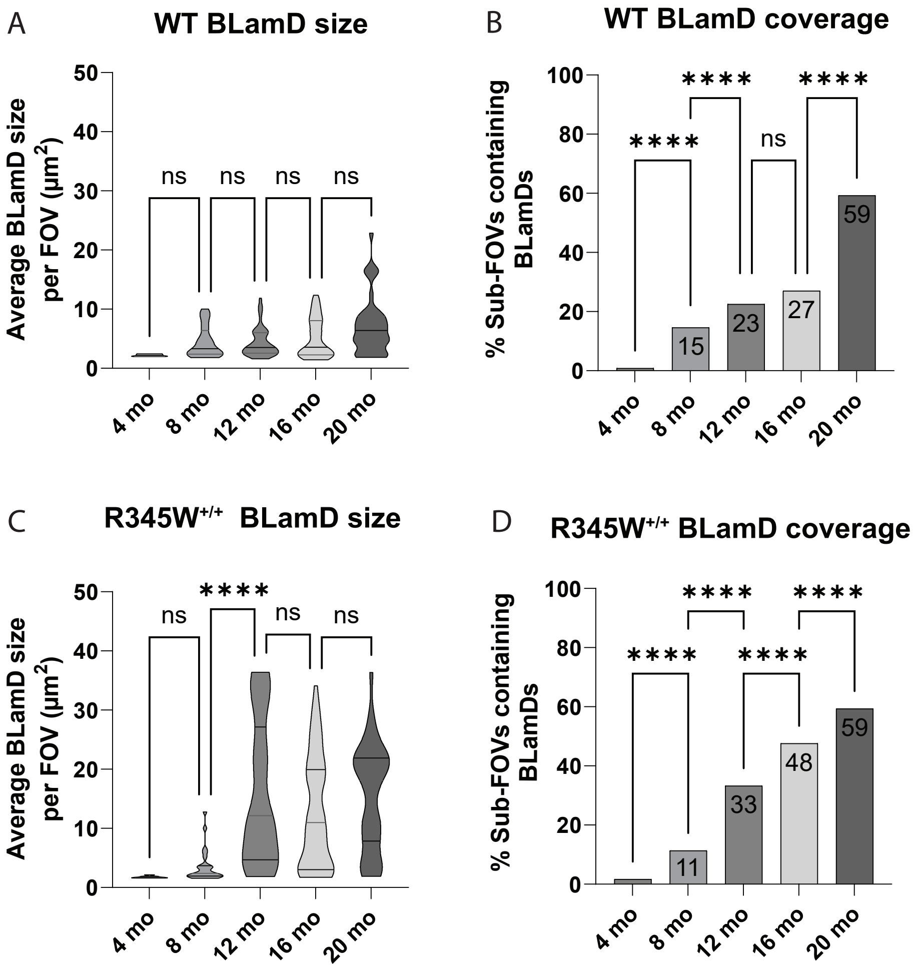

### Supplemental Fig 5

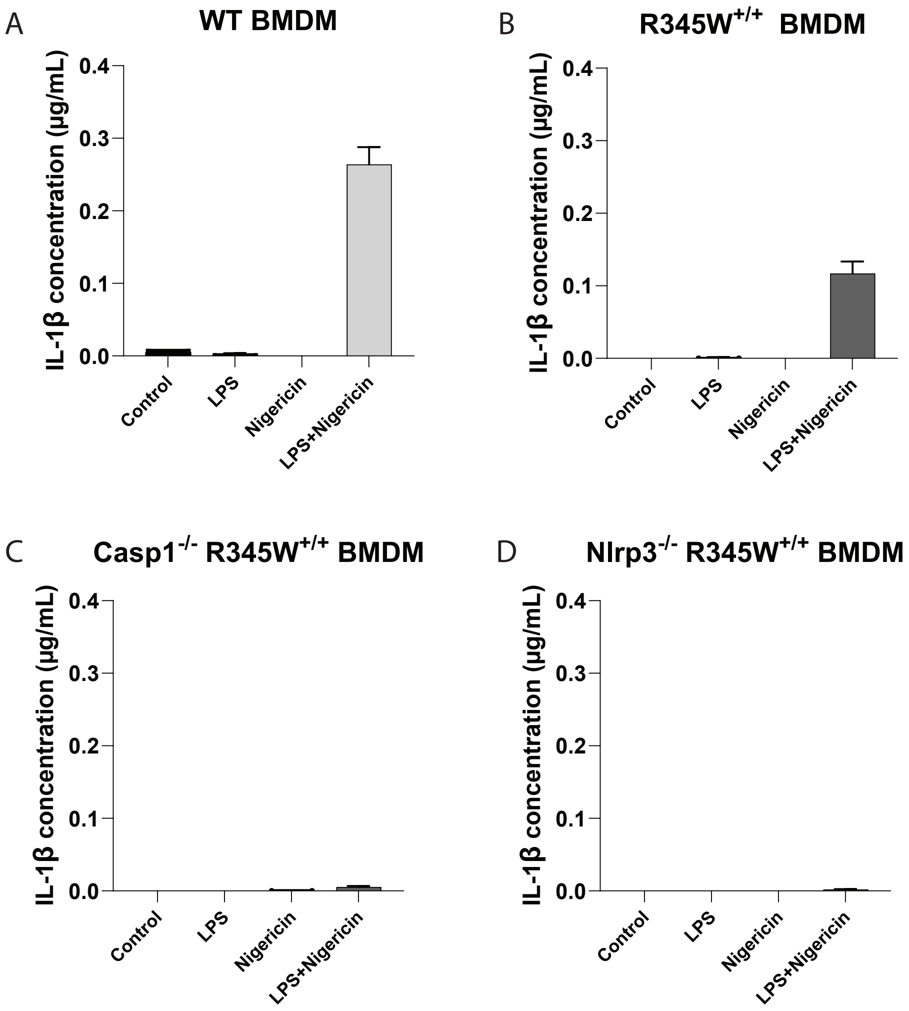

### Supplemental Fig 6

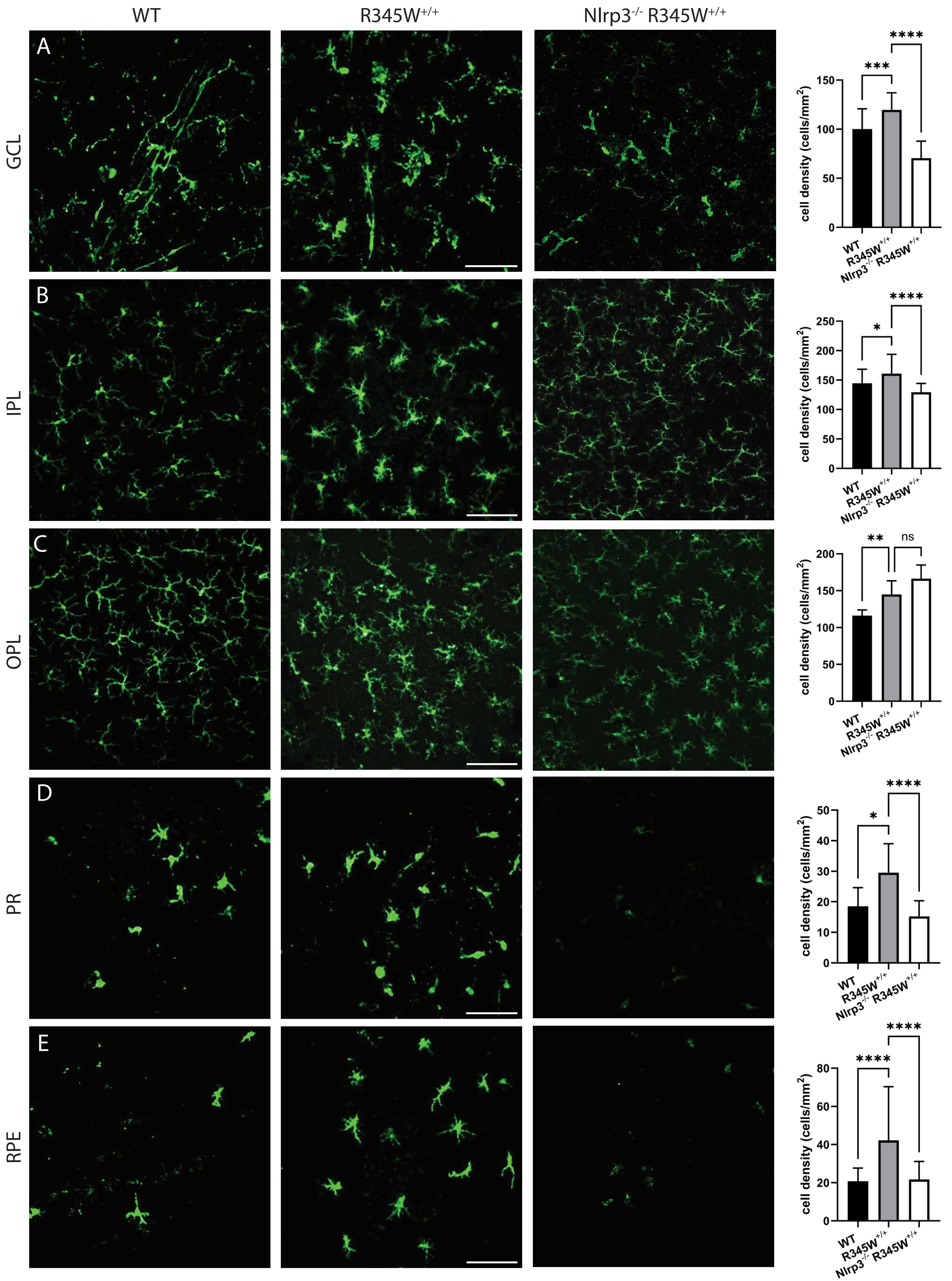

### Supplemental Fig 7

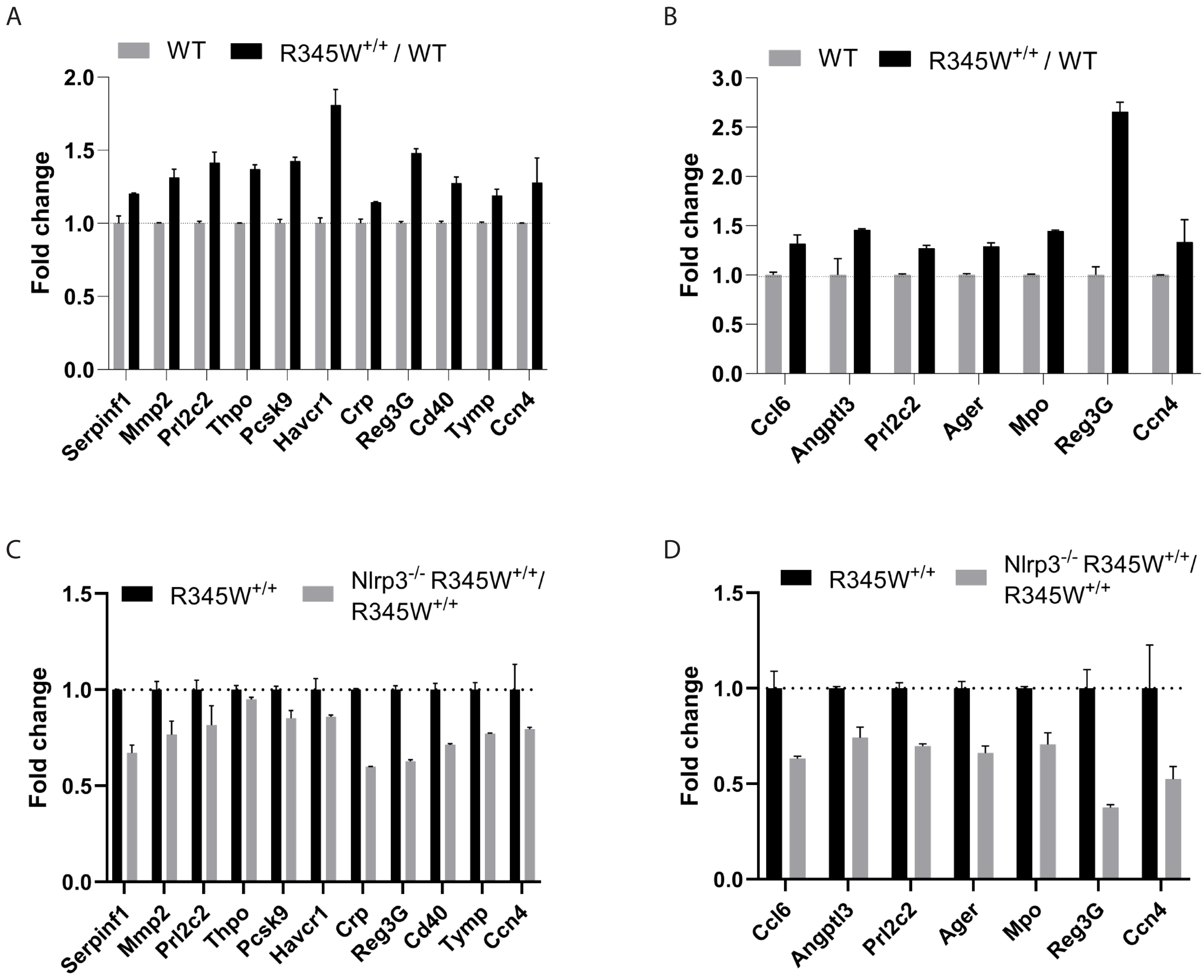
